# *STAT3* gain-of-function mutations connect leukemia with autoimmune disease by pathological dysregulation of NKG2D^hi^ CD8 killer T cells

**DOI:** 10.1101/2022.02.11.480027

**Authors:** Etienne Masle-Farquhar, Katherine J.L. Jackson, Timothy J. Peters, Ghamdan Al-Eryani, Mandeep Singh, Kathryn J. Payne, Geetha Rao, Gabrielle Apps, Jennifer Kingham, Christopher J. Jara, Ksenia Skvortsova, Alexander Swarbrick, Cindy S. Ma, Daniel Suan, Gulbu Uzel, Ignatius Chua, Jennifer W. Leiding, Kaarina Heiskanen, Kahn Preece, Leena Kainulainen, Michael O’Sullivan, Megan A. Cooper, Mikko R.J. Seppänen, Satu Mustjoki, Shannon Brothers, Tiphanie P. Vogel, Robert Brink, Stuart G. Tangye, Joanne H. Reed, Christopher C. Goodnow

## Abstract

The association between cancer and autoimmune disease is unexplained, exemplified by T-cell large granular lymphocytic leukemia (T-LGL) where gain-of-function somatic mutations in *STAT3* correlate with co-existing autoimmunity. To resolve whether these mutations are the cause or consequence of CD8 clonal expansions and autoimmunity, here we analyse patients with germline *STAT3* GOF syndrome and mice with the T-LGL mutation *STAT3^K658N^*or the most common germline mutation, *STAT3^T716M^*. STAT3 GOF mutations drove accumulation of effector CD8 T cell clones highly expressing the NKG2D receptor for MHC-I-related molecules expressed on stressed cells, and the genes for inflammatory/cytotoxic granzymes, perforin, interferon-γ and *Ccl5*/Rantes. CD8 cells were essential to lethal disease in *Stat3^K658N^*mice and their accumulation required NKG2D and the receptor for IL-15 and IL-2, IL2RB. These results demonstrate that *STAT3* GOF mutations cause effector CD8 T cell oligoclonal accumulation and that these rogue T cells contribute to autoimmune pathology, supporting the hypothesis that somatic mutations in leukemia/lymphoma driver genes contribute to autoimmune disease.

**IN BRIEF:** Leukemia and autoimmune-associated *STAT3* gain-of-function mutations dysregulate CD8 T cells to cause autoimmune pathology and oligoclonal expansion of cytotoxic killer CD8 T cells, that depend upon NKG2D and IL2RB receptors for signals displayed on stressed, damaged, infected, or mutated tissues.

## INTRODUCTION

The pathogenesis of autoimmune disease is incompletely understood. One hypothesis is that this diverse set of chronic diseases shares molecular roots with leukemia and lymphoma, wherein self-tissues are damaged by rogue clones of lymphocytes that bypass immune tolerance checkpoints through acquisition of somatic mutations in individual lymphoma/leukemia driver genes (Goodnow, 2007, Burnet, 1965).

T cell large granular lymphocytic leukemia (T-LGL) is a striking but unexplained example of the intersection between lymphoproliferative disease and autoimmunity (Lamy et al., 2017, Zhang et al., 2010). CD8 T-LGL is an indolent disorder defined by accumulation in blood of large granular CD8^+^ lymphocytes in numbers 4-10 times higher than age-matched controls, representing one or more expanded clones of terminal effector CD8 T cells with phenotype CD3^+^ TCRαβ^+^ CD8^+^ CD4^-^ CD28^-^ CD27^-^ CD45RA^+^ CD45RO^-^ CD57^+^ perforin^+^ granzyme-b^+^. T-LGL is often diagnosed because of pathological deficiency of neutrophils or erythrocytes that is presumed to reflect autoimmune destruction of these cells, and T-LGL is frequently accompanied by autoimmune rheumatoid arthritis and a diverse range of other autoimmune diseases (Lamy et al., 2017, Zhang et al., 2010). Approximately half of T-LGL patients harbour one or more expanded CD8 T cell clones bearing a gain-of-function (GOF) somatic mutation in *Signal Transducer and Activator of Transcription 3* (*STAT3*), and these patients more frequently have rheumatoid arthritis (Koskela et al., 2012). Remarkably, 8% of T-LGL patients have two, three or four independent *STAT3*-mutant CD8 T cell clones and 43% of these individuals have rheumatoid arthritis (Rajala et al., 2015), contrasting with a population frequency for rheumatoid arthritis of less than 1%. The correlation could either be an effect of autoimmune disease such as greater T cell proliferation increasing somatic mutations, or alternatively the presence of *STAT3* GOF mutations in CD8 T cells may cause autoimmune pathology.

STAT3 is a cytosolic protein that undergoes phosphorylation, dimerization and nuclear translocation to activate gene transcription in response to myriad cytokines including IL-6, IL-10, IL-15, IL-21, IL-23 and IL-27 (Deenick et al., 2018, O’Shea et al., 2013). CD8 memory T cell persistence and function depend upon STAT3, based on loss-of-function *STAT3* mutations in humans (Ives et al., 2013, Siegel et al., 2011) and mice (Cui et al., 2011). *STAT3* mutations in T-LGL are overwhelmingly missense, clustering in a small number of amino acids at the dimerization interface of the *Src* homology 2 (SH2) domain (**Figure 1A**), creating GOF substitutions that increase hydrophobicity and dimerization (Andersson et al., 2016, Barila et al., 2019, Fasan et al., 2013, Ishida et al., 2014, Jerez et al., 2012, Jerez et al., 2013, Kerr et al., 2019, Koskela et al., 2012, Morgan et al., 2017, Qiu et al., 2013, Shi et al., 2018, de Araujo et al., 2019).

**Figure 1.**
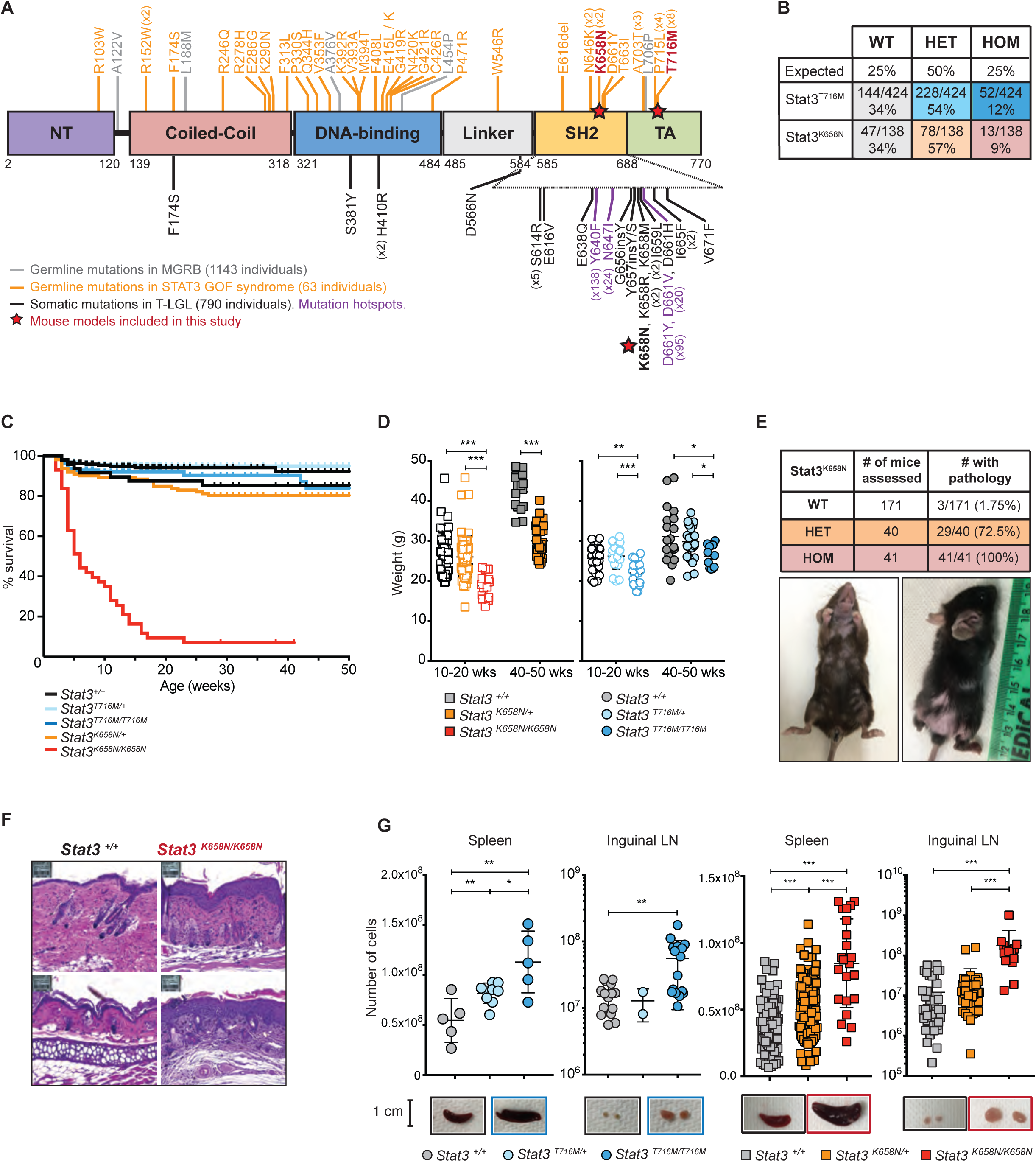
Germline *STAT3* GOF mutations cause developmental lethality, reduced weight, skin inflammation, and lymphoproliferation in mice. **A.** Schematic of STAT3 protein domains showing location of somatic mutations in T cell large granular lymphocytic leukemia (T-LGL) (Fasan et al., 2013, Jerez et al., 2012, Koskela et al., 2012, Ishida et al., 2014, Qiu et al., 2013, Barila et al., 2019, Kerr et al., 2019, Shi et al., 2018), germline mutations in STAT3 GOF syndrome and mutations identified amongst 1143 healthy individuals in the Medical Genome Reference Bank (MGRB). **B**. Expected and observed numbers and percentages of offspring of the indicated genotypes from intercrossed heterozygous parents. Statistical testing for no difference relative to an expected 1WT:2HET:1HOM ratio = p < 0.0001 for both *Stat3^T716M^*and *Stat3^K658N^*by Chi-Square test with *n* = 2 degrees of freedom. **C.** Kaplan-Meier survival curve for mice of the indicated genotypes, calculated using the product-limit method accounting for censored mice. **D.** Symbols denote weight of individual 10-20 or 40-50 week old mice of the indicated genotypes. **E.** Top, number and percentage of *Stat3^K658N^*mice that survived beyond 7 weeks of age and exhibited at least one pathology: skin inflammation, hair loss, ringtail, cataracts, diarrhoea. Bottom, representative images of hair loss and inflammation in a *Stat3^K658N/K658N^*mouse. **F.** Representative hematoxylin and eosin-stained sections of skin from the face (top) or ear pinna (bottom), from a *Stat3^+/+^*control (left) or *Stat3^K658N/K658N^* (right) mouse. **G.** Symbols denote total number of leukocytes per spleen or inguinal lymph nodes from individual mice of the indicated genotypes. Representative images of these organs shown below. **D,G.** Statistical comparisons by *t*-test corrected for multiple comparisons using the Holm-Sidak method. * *p* < 0.05; ** *p* < 0.01; *** *p* < 0.001. Data are represented as mean ± SD.

Whether *STAT3* mutations are a cause or consequence of CD8 cell clonal expansion and autoimmune disease in T-LGL is unclear because many of the mutations are in small clones representing less than 5% of CD8 cells (Rajala et al., 2015). People *without* T-LGL also have small clones of CD8 T cells with the same spectrum of somatic GOF *STAT3* mutations: in patients with Felty syndrome comprising rheumatoid arthritis, neutropenia and splenomegaly (Savola et al., 2018); in patients with aplastic anemia (Jerez et al., 2013) and primary red cell aplasia (Kawakami et al., 2018); in patients with multiple sclerosis but also in matched healthy controls (Valori et al., 2021); and in healthy blood donors with chronic HTLV-2 infection (Kim et al., 2021). Moreover, transplant of mice with bone marrow retrovirally expressing GOF mutant *STAT3* did not cause detectable CD8 T cell lymphoproliferative disease in recipient mice (Couronne et al., 2013, Dutta et al., 2018).

Compared to the SH2 domain *STAT3* somatic mutations in T-LGL, a diverse array of germline missense *STAT3* GOF mutations cause a spectrum of childhood autoimmune diseases, lymphoproliferation and immune deficiency, through mechanisms that are unclear (Flanagan et al., 2014, Haapaniemi et al., 2015, Milner et al., 2015). The most frequent clinical manifestations of germline STAT3 GOF syndrome are lymphoproliferation (diffuse lymphadenopathy, splenomegaly), autoimmune cytopenias, growth delay, enteropathy, inflammatory skin diseases, interstitial lung disease, and early-onset endocrinopathies (Fabre et al., 2019), but no consistent changes in circulating CD8 T cells have been reported. The most frequent germline mutations causing *STAT3* GOF syndrome are T716M and P715L, found in 17% and 12% of patients, respectively (Flanagan et al., 2014, Milner et al., 2015, Fabre et al., 2019). T716 and P715 are in the transactivation (TA) domain, an unstructured C-terminal segment of the protein containing the essential Y705 phosphorylation site for Janus tyrosine kinases (JAKs). (**Figure 1A**). Curiously, neither T716M nor P715L have been found as somatic mutations in 790 cases of T-LGL (Fasan et al., 2013, Jerez et al., 2012, Koskela et al., 2012, Ishida et al., 2014, Qiu et al., 2013, Barila et al., 2019, Kerr et al., 2019, Shi et al., 2018), whereas the T-LGL SH2 domain mutation K658N has been found as a germline mutation in two cases of STAT3 GOF syndrome (Flanagan et al., 2014, Ding et al., 2017). This dichotomy poses the question of whether *STAT3* GOF mutations in the SH2 and TA domains have different consequences for CD8 T cell homeostasis and autoimmunity.

Here, we analysed the consequences of *STAT3* GOF mutations in the SH2 domain (K658N) or TA domain (T716M) introduced into the mouse germline, focusing on whether or not they are sufficient to dysregulate CD8 T cells. *Stat3^K658N^*or *Stat3^T716M^*GOF mutations caused incompletely penetrant embryonic lethality, failure to thrive, inflammatory disease and lymphoproliferation – the severity of which depended on gene-dosage. Flow cytometry, deep TCR mRNA sequencing, and single-cell mRNA sequencing (scRNA-seq) analyses of *Stat3*-mutant mice and of people with germline STAT3 GOF syndrome revealed dramatic accumulation of numerous clones of CD8 T cells expressing NKG2D, the T cell and NK cell costimulatory receptor for a family of stress-induced MHC-I-like membrane proteins. These CD8 T cell clones expressed a suite of genes and proteins required for CD8 and NK cell effector and killer functions, including interferon (IFN)-γ, granzymes and perforin. CD8 T cell depletion experiments showed that these cells were required for the inflammatory wasting disease in STAT3 GOF mice, and gene targeting and receptor blockade showed that the pathological accumulation of *Stat3*-GOF NKG2D^+^ CD8 T cells depended upon NKG2D and on the IL-2Rβ receptor for IL-15, but was not diminished by individually blocking receptors for IL-6, IL-10 or IL-21. The findings demonstrate that *STAT3* GOF mutations cause T-LGL-like effector CD8 T cell oligoclonal accumulation and these rogue T cells contribute to autoimmune pathology, supporting the hypothesis that somatic mutations in leukemia/lymphoma driver genes contribute to autoimmune disease.

## RESULTS

### Germline gain-of-function *Stat3* mutations cause pathology in mice

Two *STAT3* GOF mutations were selected for investigation in CRISPR/*Cas9* genome-edited mice, based on the pattern of somatic and germline *STAT3* GOF mutations in humans (**Figure 1A**).

K658N, a somatic mutation in a patient with T-LGL and pancytopenia (Koskela et al., 2012), was chosen to model somatic mutations in the SH2 domain dimerization interface that dominate T-LGL. Importantly, the same mutation has been detected in the germline in two unrelated children with lymphadenopathy, splenomegaly, autoimmune cytopenia, dermatitis, recurrent upper respiratory infections, autoimmune enteropathy, small stature, and delayed puberty (Flanagan et al., 2014, Ding et al., 2017). The second mutation, T716M in the TA domain, was chosen because it is the most frequent germline mutation causing STAT3 GOF syndrome. We had no difficulty generating T716M heterozygous founder mice in the homogeneous C57BL/6J genetic background. Of 279 microinjected C57BL/6 zygotes, 13 pups were born and 2 had a homologously-recombined T716M allele. By contrast, we were unable to generate K658N founders in the inbred C57BL/6 background. Of 266 microinjected C57BL/6 zygotes, 9 pups were born but none carried the homologously-recombined K658N allele. However, when (C57BL/6J x FVB/J) F_2_ hybrid zygotes were microinjected with the same sgRNA and targeting oligonucleotide, 19 pups were born from 129 injected zygotes and 2 had homologously-recombined K658N alleles. This result could reflect random Poisson sampling among small numbers of pups, or it may reflect a need for hybrid vigour to obtain viable K658N heterozygous animals.

Following intercross of *Stat3^T716M/+^* or *Stat3^K658N/+^* heterozygotes, homozygotes were observed at less than half the expected Mendelian frequency at time of weaning (**Figure 1B**). In addition, 50% of *Stat3^K658N/K658N^*homozygotes died or required ethical culling by age 7 weeks (**Figure 1C**). At 10-20 and 40-50 weeks of age, *Stat3^T716M^*and *Stat3^K658N^*mutant mice weighed less than wild-type littermate controls (**Figure 1D**).

From 3-5 weeks of age, *Stat3^K658N/K658N^* homozygotes were reduced in size, hunched, with reduced movement and skin inflammation resulting in red, flaky ears and variable hair loss around the eyes, on the face, snout, neck and tail (**Figure 1E**). At lower penetrance, they presented with cataracts, ringtail, red and inflamed joints. This visible, gross pathology was fully penetrant in *Stat3^K658N/K658N^* mice that survived beyond 7 weeks, and though milder, eye and skin pathology appeared on average at 15 weeks of age in 75% of *Stat3^K658N/+^*mice (**Figure 1E**). *Stat3^T716M/T716M^*homozygous mice presented at 7-25 weeks of age with cataracts and skin inflammation limited to the ears and tail – a phenotype that occurred at low penetrance in *Stat3^T716M/+^*mice at 40-50 weeks of age.

Comprehensive anatomical and histopathology analysis of *Stat3^T716M^*homozygous (*n*=3) and wild-type littermates (*n*=3), and *Stat3^K658N^*homozygous (*n*=5), heterozygous (*n*=1) and wild-type littermates (*n*=5) revealed consistent abnormalities in *Stat3* mutants were primarily in keratinized epithelia. In the skin, 5/5 *Stat3^K658N/K658N^*mice had acanthotic epidermal hyperplasia and dermal infiltrates of mononuclear cells and well-differentiated mast cells especially on the face and ears (**Figure 1F**), tail, penis in males, and milder dermatitis of distal limbs. 5/5 mice had inflammation of the middle ear with submucosal oedema and large foamy luminal macrophages, and 3/5 displayed acanthosis and lymphocytic or neutrophilic infiltration of the esophageal mucosa.

Splenomegaly and lymphadenopathy – the two most prevalent abnormalities in human germline STAT GOF syndrome - were highly penetrant along with blood leukocytosis in mice homozygous for either the K658N or T716M mutations, while heterozygotes had intermediate increases in spleen and blood leukocytes (**Figure 1G**, **Supplementary Figure 1A**).

Collectively, these results indicate that both of the *STAT3* GOF mutations analysed were sufficient to cause lymphadenopathy accompanied by inflammatory skin disease, and partial lethality during development, with these measures tending to be more severe for mice with the K658N mutation in the SH2 domain than mice with the T716M mutation in the TA domain. The differences in gross pathology between the two strains may reflect a functional difference between the two GOF alleles, but is equally likely to reflect differences in other genes between C57BL/6 and FVB mouse strains which profoundly affect traits such as skin keratinocyte transformation (Wakabayashi et al., 2007).

### Germline STAT3 gain-of-function causes accumulation of NKG2D^+^ effector CD8 T cells in mice

We next focused on the question of whether *Stat3* GOF mutations are sufficient alone to dysregulate CD8 T cells. *Stat3^T716M/T716M^* and *Stat3^K658N/K658N^* mice had increased frequencies of peripheral blood CD8^+^ relative to CD4^+^ T cells (**Supplementary Figure 1B**), and mildly increased total number of CD8 T cells per spleen (**Figure 2A**). However, *Stat3^K658N/K658N^*and *Stat3^T716M/T716M^*mice had a dramatic, 8- or 7-fold respective increase in percentage and 13- or 12-fold respective increase in number of CD8 T cells with a CD62L^-^ CD44^+^ effector memory phenotype (**Figure 2B,C**). *Stat3^K658N/+^*and *Stat3^T716M/+^*mice also had an intermediate 6- or 2-fold respective increase in number of effector memory CD8 T cells (**Figure 2B,C**). Thus, both GOF mutations significantly dysregulate CD8 T cells.

**Figure 2.**
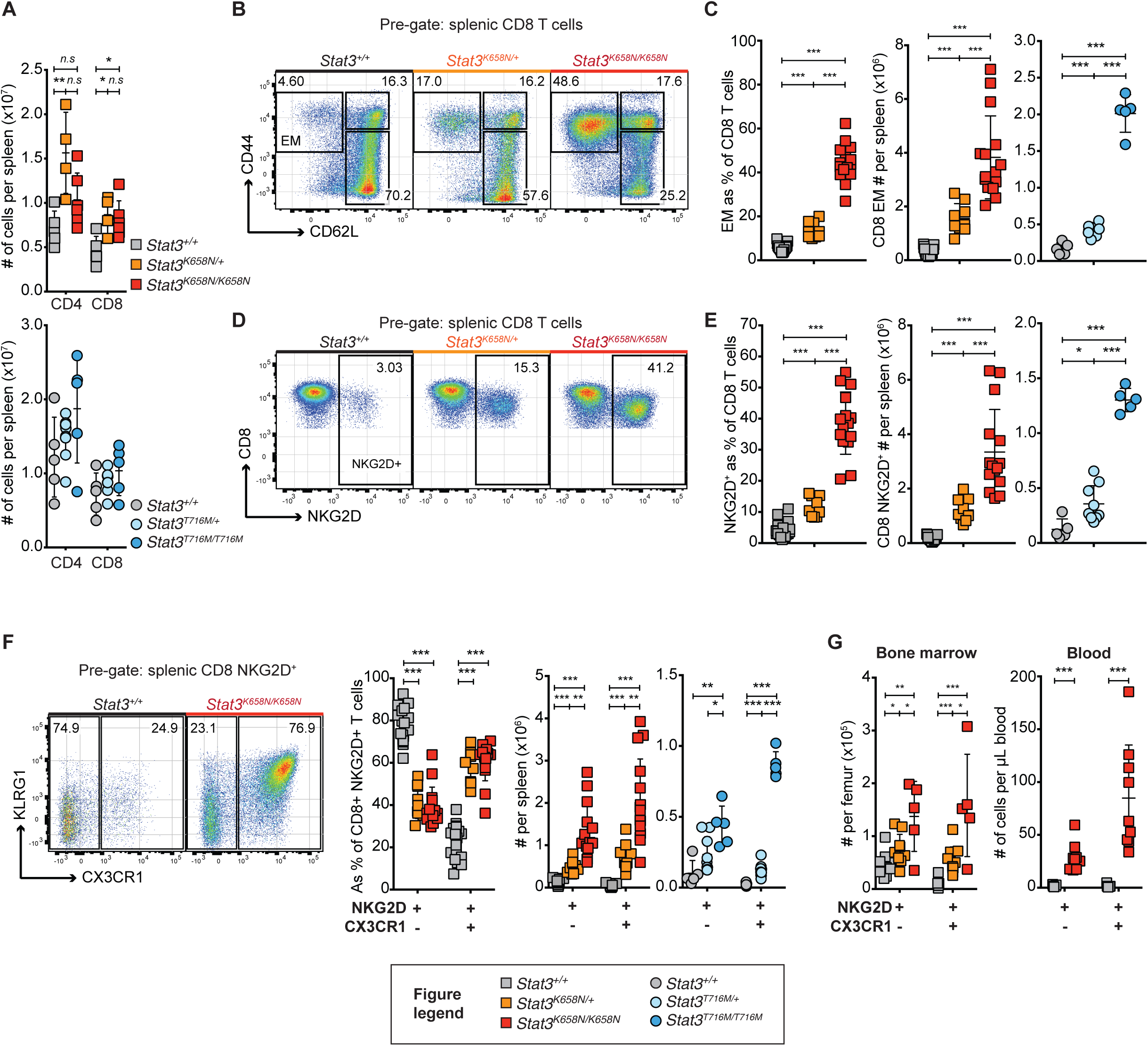
Germline *Stat3* GOF mutations drive accumulation of NKG2D^+^ CX3CR1^+^ KLRG1^high^ CD8 T cells. **A.** Symbols show total number of CD4^+^ and CD8^+^ T cells per spleen in individual mice of the indicated genotypes. **B.** Representative flow cytometric analysis of spleen cells from mice of the indicated genotypes gated on CD8 T cells, showing the percentage in CD44^-^ CD62L^+^ naïve, CD44^+^ CD62L^+^ Central memory (CM), and CD44^+^ CD62L^-^ effector memory (EM) subsets. **C.** Symbols show percentage of CD8 T cells with an EM phenotype, and total number per spleen of EM CD8 T cells in individual mice of the indicated genotypes. **D.** Representative flow cytometric analysis of C-type lectin-like receptor NKG2D expression by splenic CD8 T cells. **E.** Percentage of CD8 T cells expressing NKG2D, and total number per spleen of NKG2D^+^ CD8 T cells. **F.** Left, representative flow cytometric analysis of CX3CR1 and KLRG1 expression by *Stat3^+/+^* (black) or *Stat3^K658N/K658N^*CD8 NKG2D^+^ T cells. Right, percentage and total number per spleen of CX3CR1^-^ NKG2D^+^ or CX3CR1^+^ NKG2D^+^ CD8 T cells in individual mice of the indicated genotypes. **G.** Total number of CX3CR1^-^ or CX3CR1^+^ NKG2D^+^ CD8 T cells, per hind leg femur (left) or per Lμ blood (right) in individual mice. **C-G**, genotypes as indicated in A and key. Statistical comparisons made by *t*-test, corrected for multiple comparisons using the Holm-Sidak method. * *p* < 0.05; ** *p* < 0.01; *** *p* < 0.001. Data are representative of *n* > 5 independent experiments with *n* > 4 mice per group. Data are represented as mean ± SD.

To analyse effector CD8 T cell dysregulation further, we focussed initially on the expression of NKG2D for three reasons. First, *Klrk1* (encoding NKG2D) is a STAT3 target gene in NK cells (Zhu et al., 2014), and acts as an activating receptor for the stress-induced MHC class I-related proteins MICA and ULBP in humans (Bauer et al., 1999, Wu et al., 1999, Sutherland et al., 2001) and Rae1 and H-60 in mice (Cerwenka et al., 2000, Diefenbach et al., 2000). Second, most CD8^+^ CD57^+^ T-LGLs express NKG2D (Bigouret et al., 2003). Third, effector memory CD8 T cells expressing NKG2D cause autoimmune alopecia in C3H/H3J mice, a condition treated in mice and humans by JAK-STAT inhibition (Xing et al., 2014). In mice, NKG2D is expressed upon CD8 T cell activation (Diefenbach et al., 2000, Jamieson et al., 2002). We observed a *Stat3* mutant allele dose-dependent increase in the percentages of CD44^+^ CD62L^-^ effector memory CD8 T cells expressing NKG2D in the blood, bone marrow and spleen of *Stat3^K658N^*and *Stat3^T716M^*mutant mice (**Supplementary Figure 1C**). Increased numbers of CD8^+^ NKG2D^+^ T cells in these mice are therefore not explained by increased numbers of effector memory CD8 T cells. Strikingly, *Stat3^K658N/K658N^*and *Stat3^T716M/T716M^*mice had a 9- or 7-fold increase in percentage and 20- or 12-fold increase in number, respectively, of NKG2D^+^ CD8^+^ T cells in the spleen, relative to wild-type controls (**Figure 2D,E**). *Stat3^K658N/+^*and *Stat3^T716M/+^*heterozygotes had intermediate 8- or 3-fold respective increases in number of splenic NKG2D^+^ CD8^+^ T cells (**Figure 2D,E**).

To further characterise this expanded population of effector CD8^+^ T cells, we measured other markers of migration, maturation and activation. In wild-type mice, only 20% of CD8 NKG2D^+^ T cells expressed the atypical chemokine receptor CX3CR1, whereas this subset increased to 60% of CD8 NKG2D^+^ T cells in heterozygous or homozygous mutant mice from either *Stat3* GOF strain (**Figure 2F**). This resulted in a particular expansion of CD8^+^ NKG2D^+^ CX3CR1^+^(and to a lesser extent CX3CR1^-^) T cells in the spleen, bone marrow and blood of *Stat3^K658N^*and *Stat3^T716M^*mutant mice relative to wild-type controls (**Figure 2F, G**). CD8^+^ NKG2D^+^ CX3CR1^+^ T cells in both *Stat3*-wild-type and -mutant mice exhibited a TCRβ CD44^+^ CD62L^-^ KLRG1^high^ CD27^low^ CD127^low^ and CXCR3^low^phenotype of terminally differentiated effector CD8 T cells (**Figure 2F**, **Supplementary Figure 1D**). NKG2D^+^ CX3CR1^+^ CD8 T cells expressed the lowest cell-surface levels of TCRβ CD8 among CD8 T cell subsets, and this was further diminished in *Stat3*-mutant mice and (**Supplementary Figure 1E**), reminiscent of CD8 and TCR “de-tuning” during CD8 cytotoxic responses (Xiao et al., 2007). Collectively, these results reveal that two independent *Stat3* GOF mutations drive aberrant accumulation of NKG2D^+^ effector memory CD8 T cells in mice.

### Germline STAT3 gain-of-function causes accumulation of NKG2D^hi^and CD57^+^ effector CD8^+^ T cells in humans

To generalize the findings in *Stat3*^GOF^mice, we investigated the peripheral blood CD8^+^ T cell compartment in 11 people with childhood-onset multi-organ autoimmunity and lymphoproliferation caused by germline *STAT3* GOF mutations (listed in **Supplementary Table 1**). The detailed clinical features of the patients will be described elsewhere (Leiding *et al*., in preparation).

Flow cytometric analysis of peripheral blood mononuclear cells (PBMCs) revealed a normal frequency of circulating T cells (**Figure 3A**). However, similar to *Stat3*-GOF mutant mice, we observed a significant relative increase in CD8 and decrease in CD4 T cell frequency, with a consequent 2-fold decrease in CD4:CD8 ratio, in patients relative to controls (**Figure 3B**). There were no significant differences in percentages of CD8 T cells with a CD45RA^+^ CCR7^+^ naïve, CD45RA^-^ CCR7^+^ Central memory, CD45RA^-^ CCR7^-^ effector memory or CD45RA^+^ CCR7^-^ terminal effector memory re-expressing CD45RA (T_EMRA_) phenotype between STAT3 GOF patients and healthy donors (**Figure 3C**). While all CD8 T cells express NKG2D in humans, we observed a large increase in the mean fluorescence intensity of NKG2D on patient CD8 T cells, particularly on the T_EMRA_ subset, compared to healthy donors analysed in parallel (**Figure 3D**; **Supplementary Figure 2A,B**). Consequently, patients had a significant increase in NKG2D^hi^cells as a percentage of CD8 T cells, T cells or of circulating lymphocytes (**Figure 3E**, **Supplementary Figure 2C**). Patients also had a 3-fold increase in the mean frequency of CX3CR1^+^ NKG2D^hi^CD8 T cells as a percentage of CD8 T cells or of circulating lymphocytes (**Figure 3F**). Consistent with T_EMRA_ and T-LGL cells, CD8 T cells from *STAT3^GOF^*patients P9 and P3 expressed lower levels of CD28, CD127, CD27, CCR7, and higher CD95, relative to healthy controls (**Supplementary Figure 3A**).

**Figure 3.**
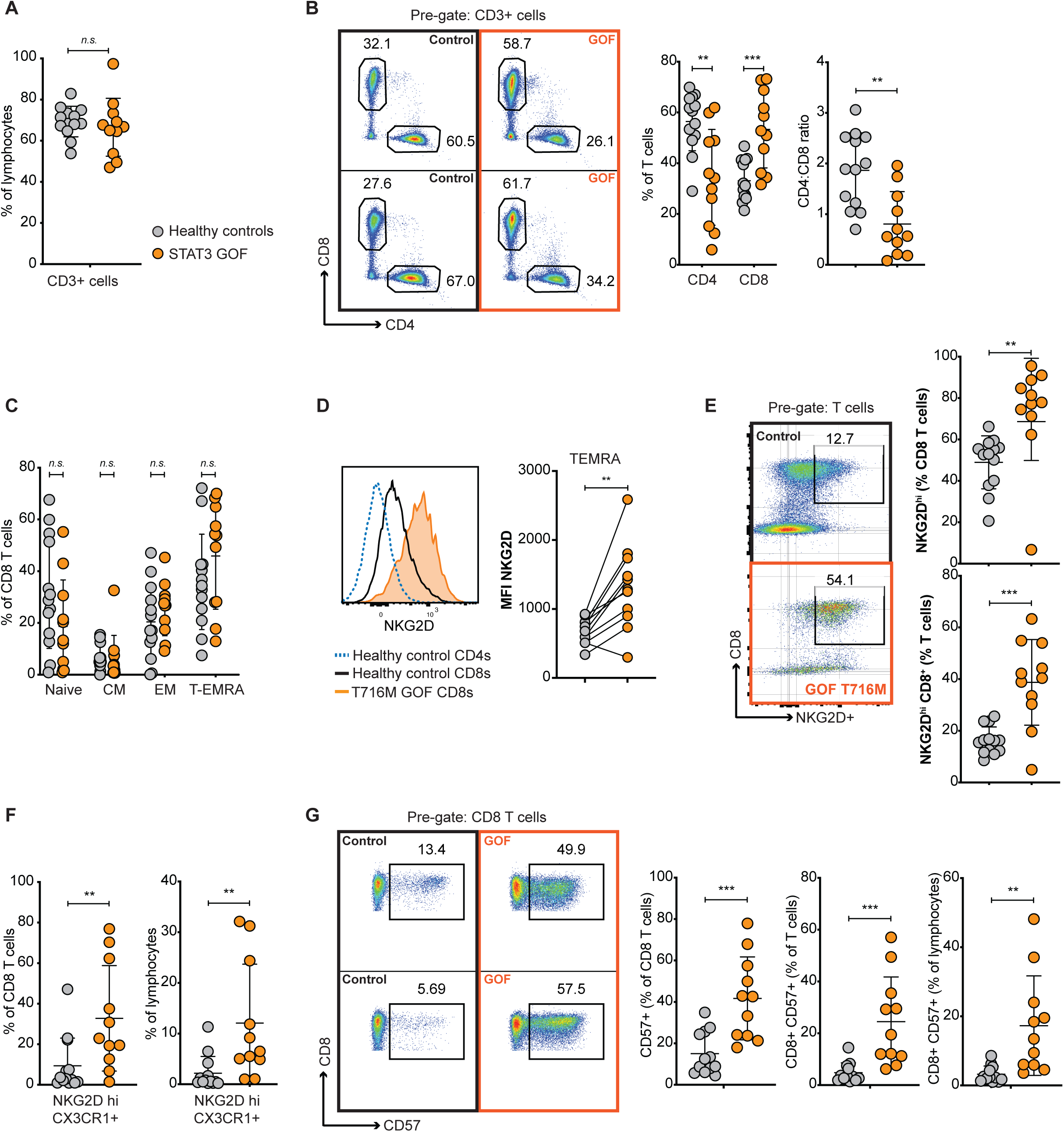
NKG2D^hi^or CD57^+^ effector CD8 T cells accumulate in individuals with *STAT3* GOF syndrome. **A-F.** Flow cytometric analysis of peripheral blood T cells in 13 healthy donors (grey) and 11 patients with germline *STAT3* GOF syndrome (orange). **A.** Percentage of circulating lymphocytes expressing CD3. **B.** Representative flow cytometric analysis (in two patients and two healthy donors) and percentages of CD4^+^ or CD8^+^ T cells and resulting CD4:CD8 ratio. **C.** Percentages of CD8 T cells with a CD45RA^+^ CCR7^+^ naïve, CD45RA^-^ CCR7^+^ Central memory, CD45RA^-^ CCR7^-^ effector memory and CD45RA^+^ CCR7^-^ terminal effector re-expressing CD45RA (T_EMRA_) phenotype. **D.** Left, Representative flow cytometric histograms showing distribution of NKG2D fluorescence on patient and healthy control CD8 T cells and control CD4 cells. Right, NKG2D mean fluorescence intensity (MFI) on T_EMRA_ CD8 T cells in individual patients and controls. Lines join samples analysed in a same experiment. In experiments that included multiple healthy controls, grey symbols denote the average of the healthy control MFIs. **E.** Representative plots of NKG2D and CD8 fluorescence on patient and healthy control T cells, and percentage of NKG2D^hi^CD8^+^ T cells among CD8 T cells (top) or total T cells (bottom). **F**. CX3CR1^+^ NKG2D^hi^CD8+ T cells, as a percentage of CD8^+^ T cells or of total lymphocytes. **G.** Representative flow cytometric analysis and frequency of CD57^+^ CD8^+^ T cells, as a percentage of CD8 T cells, total T cells or total lymphocytes. **B-G**, cohort as indicated in A and key. Data are combined from *n* = 9 independent experiments. Statistical comparisons made by *t*-test, corrected for multiple comparisons using the Holm-Sidak method. * *p* < 0.05; ** *p* < 0.01; *** *p* < 0.001. Data are represented as mean ± SD.

Since the majority of T-LGL and a small subset of granzyme^hi^perforin^hi^normal CD8 T cells express the sulphated trisaccharide epitope CD57 (also called HNK-1; (Focosi et al., 2010, Chou et al., 1986, Voshol et al., 1996)), e measured CD57 expression on CD8 T cells in germline *STAT3*w GOF patients. Relative to healthy donors, patients had a 3- and 12-fold increase in CD57^+^ CD8 T cells as a percentage of CD8 T cells or of circulating lymphocytes, respectively (**Figure 3G**). In 2 out of the 8 patients where these data were able to be calculated, the total number of CD57^+^ CD8^+^ Cells was greater than 0.5 x 10^9^/L (**Supplementary Table 1**), representing the threshold for T-LGL diagnosis when accompanied by autoimmune conditions. In a third patient, the total number was 0.46 x 10^9^/L (**Supplementary Table 1**). In this limited series of patients there was no discernible correlation between the location of the mutations in different STAT3 functional domains and the degree of CD57^+^ CD8 T cell dysregulation (**Supplementary Figure 3B**). Thus, germline *STAT3* GOF mutations both within and external to the SH2 domain are sufficient to cause CD57^+^ CD8 T cell accumulation in humans.

To explore the dysregulated effector/memory CD8 T cells in human STAT3 GOF syndrome, we performed single-cell RNA and surface glycoprotein analysis of thousands of non-naïve (excluding CD45RA^+^ CCR7^+^) CD8 T cells sorted by FACS (**Figure 4A**), along with CD8-CD4+ cells as a comparator, from PBMCs of 2 healthy donors and 2 individuals with STAT3 GOF syndrome: P10 with a T716M mutation and P9 with a T663I SH2 domain mutation. CD4 T cells were excluded from further analyses, and all CD8 T cells were analysed by Receptor And Gene Expression RNA sequencing (RAGE-seq) (Singh et al., 2019)), which couples T cell receptor (TCR) mRNA long-read sequencing with global transcriptome short-read single-cell 3’ mRNA sequencing (scRNA-seq), accompanied by surface glycoprotein expression measured by staining with oligonucleotide-conjugated antibodies and Cellular Indexing of Transcriptomes and Epitopes by sequencing (CITE-seq) (Stoeckius et al., 2017).

**Figure 4.**
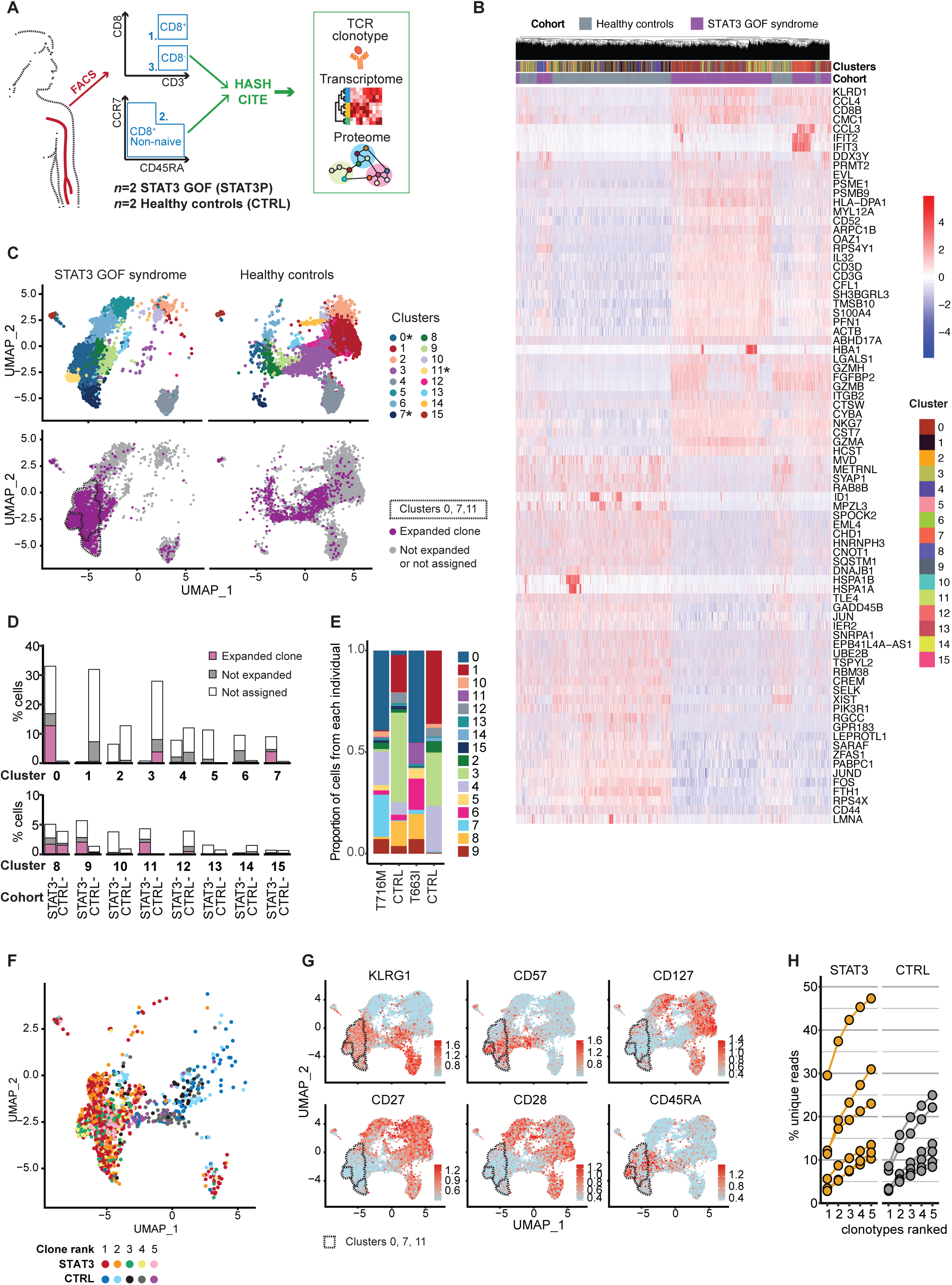
Clonally expanded CD57^+^ effector CD8 T cells expressing numerous killer cell genes in STAT3 GOF syndrome patients. **A.** Schematic workflow for analysing single non-naïve CD8 T cells from 2 healthy controls and 2 STAT3 GOF patients, P10 (T716M mutation) and P9 (T663I mutation), by sequencing of T cell receptor (TCR) mRNA clonotypes, global 3’ mRNA transcriptome, and CITE-seq tagging of cell surface proteins with oligonucleotide-conjugated antibodies. **B.** Unsupervised clustered heatmap of non-naïve CD8 cells (columns) from patients and controls (denoted by Cohort key) and relative expression (heat key, log2) of the top 40 increased and decreased genes (rows) in patient versus control non-naïve CD8**^+^**T cells. **C.** Uniform manifold approximation and projections (UMAPs) following dimensionality reduction analysis of gene expression. Top: each cell from patients or controls is denoted by a dot, colour-coded into one of sixteen cell subsets numbered 0-15 identified by graph-based clustering using K nearest neighbour (KNN; Cluster key). Bottom: cells that belong to expanded clones based on TCR V(D)J clonotypic sequence are coloured. **D.** Percentage of non-naïve CD8 cells in patients (STAT3) or controls (CTRL) corresponding to cell clusters 0-15 generated by dimensionality reduction analysis, and the fraction in each cluster corresponding to an expanded clone (≥5 cells assigned the same clonotype call based on V gene, J gene and CDR3 amino acid sequence for TRB and TRA). **E.** Relative proportions of non-naïve CD8 T cells in clusters 0-15 in the individual patients and controls. **F.** UMAP with colour coding of individual cells belonging to the five most frequent TCR clones, ranked 1-5, within patients (STAT3) or controls (CTRL) as indicated in the key. **G.** UMAPs displaying CITE-seq analysis of relative expression of the indicated cell surface markers on individual non-naïve CD8 T cells. Dashed outlines denote cell clusters 5, 0, 11 and 7, from top to bottom. **H.** Percentage of unique transcripts accounted for by the top 5 largest clonotypes in healthy controls (*n*=6) or STAT3 GOF patients (*n*=6), following deep *TCRB* VDJ mRNA sequencing of sorted NKG2D^hi^CD8 T cells. Wilcoxon rank sum test for healthy control mean being lower than STAT3 GOF patient mean: p = 0.155. Wilcoxon rank sum test for mean of control being lower than mean of STAT3, p = 0.197

Global scRNAseq profiling showed that non-naive CD8 T cells from the patients and healthy donors were transcriptionally distinct (**Figure 4B**). *STAT3*^GOF^ cells had significantly increased expression of 768 genes and decreased expression of 396 genes, relative to healthy individuals (**Supplementary Table 2**). Among the top 40 most increased were genes crucial to CD8 T cell effector functions and cytotoxicity (**Figure 4B**): *GZMA, GZMB, GZMH* encode granzymes*; GNLY* encodes the pore-forming protein granulysin; *NKG7* encodes a key regulator of cytotoxic granule exocytosis (Ng et al., 2020)*; ARPC1B* encodes a subunit of the actin-related protein 2/3 complex required for CD8 T cell survival and cytotoxicity (Randzavola et al., 2019); *PFN1* encodes perforin and *FGFBP2*-encodes the cytotoxic lymphocyte-specific protein co-expressed with perforin (Ogawa et al., 2001); *HCST* encodes DAP10, the transmembrane adaptor protein for NKG2D that stabilises NKG2D expression (Park et al., 2011) and allows signal transduction upon binding to NKG2D ligands (Wu et al., 1999). *STAT3*^GOF^CD8 T cells also had increased expression of *CCL3, CCL4, CCL5, IL-32, KLRD1, KLRG1, KLRF1, CD3G, CD8B, CD3D, CD3E, CD8A* and *LCK* (Supplementary Table 2).

Unsupervised dimensionality reduction analysis of RNA-seq transcriptome data for *STAT3* GOF and control non-naïve CD8 T cells and CD4 T cells identified 16 distinct non-naïve CD8 T cell clusters (numbered 0-15) (**Figure 4C**). Clusters #0, #7 and #11 were >90% constituted of STAT3 GOF patient cells, and RAGE-seq revealed most of the T cells in these clusters that had an assignable TCR V(D)J clonotypic sequence were members of expanded T cell clones defined as 5 or more cells sharing the same TCR V(D)J clonotypic sequence (**Figure 4C,D**). Notably, a significantly larger proportion of patient CD8 T cells were in expanded clonotypes relative to controls (60.9% versus 30.3%; 2x2 Fisher exact *p* = 0.046), and expanded clones were larger on average in the patients relative to healthy donors: 36.4 cells versus 14.3 cells, respectively (t-test *p* = 0.026).

Cluster #0 was more frequent in both patients relative to controls, whereas cluster #11 appeared unique to P9 who harbours a T663I mutation in the SH2 domain and cluster #7 was predominantly from P10 with the T716M TA domain mutation (**Figure 4E**). Clusters 0, 7 and 11 were enriched for the largest T cell clones in STAT3 GOF patients (**Figure 4F**) and expressed high levels of mRNAs required for cytotoxic effector functions (*GZMA, GZMB, GZMH*, *PFN1, FGFBP2, NKG7, CTSW, GNLY, HCST, EFHD2*), inflammatory chemokines and cytokines (*CCL3*, *CCL4, CCL5, IL32; IFNG*), and encoding TCR/co-receptor subunits (*CD8A, CD8B, CD3G, CD3D, CD3E*). In contrast, mRNAs involved in JUN/FOS (*FOS, JUNB, JUN, JUND, FOSB*) and NF-κB signalling (*LTB, TNFAIP3, CD69, NFKBIA, NFKB1, NFKB2, NFKBIZ*), as well as *SOCS3* and *SOCS1*, were reduced in these clusters (**Supplementary Table 3**).

CITE-seq surface protein expression features of cluster #0, #7 and #11 were distinguished by high levels of CD57 and CD45RA and low levels of CD55, CD28, CD27, and CD62L (**Figure 4G**; **Supplementary Table 3**). Collectively, these results demonstrate that a distinct subset of clonally expanded killer effector CD8 T cells is pathologically increased in human STAT3 gain of function syndrome.

Given the variable frequency of *STAT3* GOF mutant CD8 cell clones in human T-LGL and autoimmune disorders, we explored the range of frequencies represented by clonally expanded T cells within the NKG2D^hi^CD8 T cells from germline STAT3 GOF patients by performing deep *TCRB* VDJ mRNA sequencing on sorted NKG2D^hi^CD8 T cells from 6 patients and 6 controls (**Figure 4H**). The largest clonotypes in the STAT3 GOF patients accounted for 2.8-29.5% of unique transcripts (mean 10.80%) and the largest clonotypes in healthy controls comprised 2.8-8.5% of unique transcripts (mean 4.7%). The five largest clonotypes accounted for 10.3-47.3% of unique transcripts (mean 22.8%) in patients and 8.4-24.9% in controls (mean 15.1%). Thus, large CD8 T cell clonal expansions occur in germline *STAT3* GOF patients and in healthy controls.

### Accumulating NKG2D^+^ CD8 T cells in *Stat3* GOF mice over-express cell cycle and killer cell genes

To complement the human analysis above, single cell RNA-seq of STAT3 GOF CD8 T cells was performed on thousands of NKG2D^+^ CX3CR1^+^ CD8 T cells flow sorted from the spleen of *Stat3^T716M/T716M^*, *Stat3^K658N/K658N^*and littermate *Stat3^+/+^* mice (**Figure 5A**). For comparison, total CD8 T cells were also sorted from *Stat3^T716M/T716M^*and littermate *Stat3^+/+^*mice. During UMAP analysis, cells within the total CD8 T cell population that clustered with sorted NKG2D^+^ CX3CR1^+^ CD8 cells were excluded prior to differential gene expression analysis. In *Stat3^T716M/T716M^*mice, NKG2D^+^ CX3CR1^+^ CD8 T cells differentially expressed 4667 genes compared to total CD8 cells from the same mice (**Figure 5B**; **Supplementary Table 4**). RNAs encoding the two markers used for sorting, *Klrk1* and *Cx3cr1*, were the first and third most increased in the sorted cells, validating the analysis. Among the other significantly increased mRNAs in NKG2D^+^ CX3CR1^+^ CD8 cells were those encoding proteins mediating CD8 T cell cytotoxic and inflammatory functions, including *Gzmb, Gzmk, Gzma, Gzmm, Prf1, Ifng, Ccl5, Ccl4, Ccl3, Ccl9, Ccl6, Ctla2a, Ctla2b, Cxcr3*, and an extended suite of genes encoding NK cell-associated molecules including *Hcst* (encoding the NKG2D signalling partner DAP10), *Klrc1, Klre1, Klrc2, Klrb1c, Klra9, Klrb1f, Klra3, Klra8, Klri2, Klrd1, Klrb1a* (**Figure 5B**; **Supplementary Table 4**). CD8 memory/effector regulating genes, *Ccr7, Ccr9, Sell* (CD62L), *Il7r* (CD127), *Cd27,* and *Id3* mRNAs were decreased while *Zeb2, Prdm1, Id2, Eomes* and *Stat4* mRNA levels were increased in NKG2D^+^ CX3CR1^+^ relative to total CD8 T cells (**Figure 5B**; **Supplementary Table 4**). *Zeb2* contributes to memory/effector CD8 T cell differentiation and responses (Omilusik et al., 2015), as do *Id2*, *Id3*, *Prdm1* (encodes BLIMP-1), *Eomes* and *Stat4* (Kaech and Cui, 2012). The global scRNAseq analysis indicates that the dysregulated population of effector memory CD8 cells in STAT3 GOF mice display a gene expression profile of killer effector cells.

**Figure 5.**
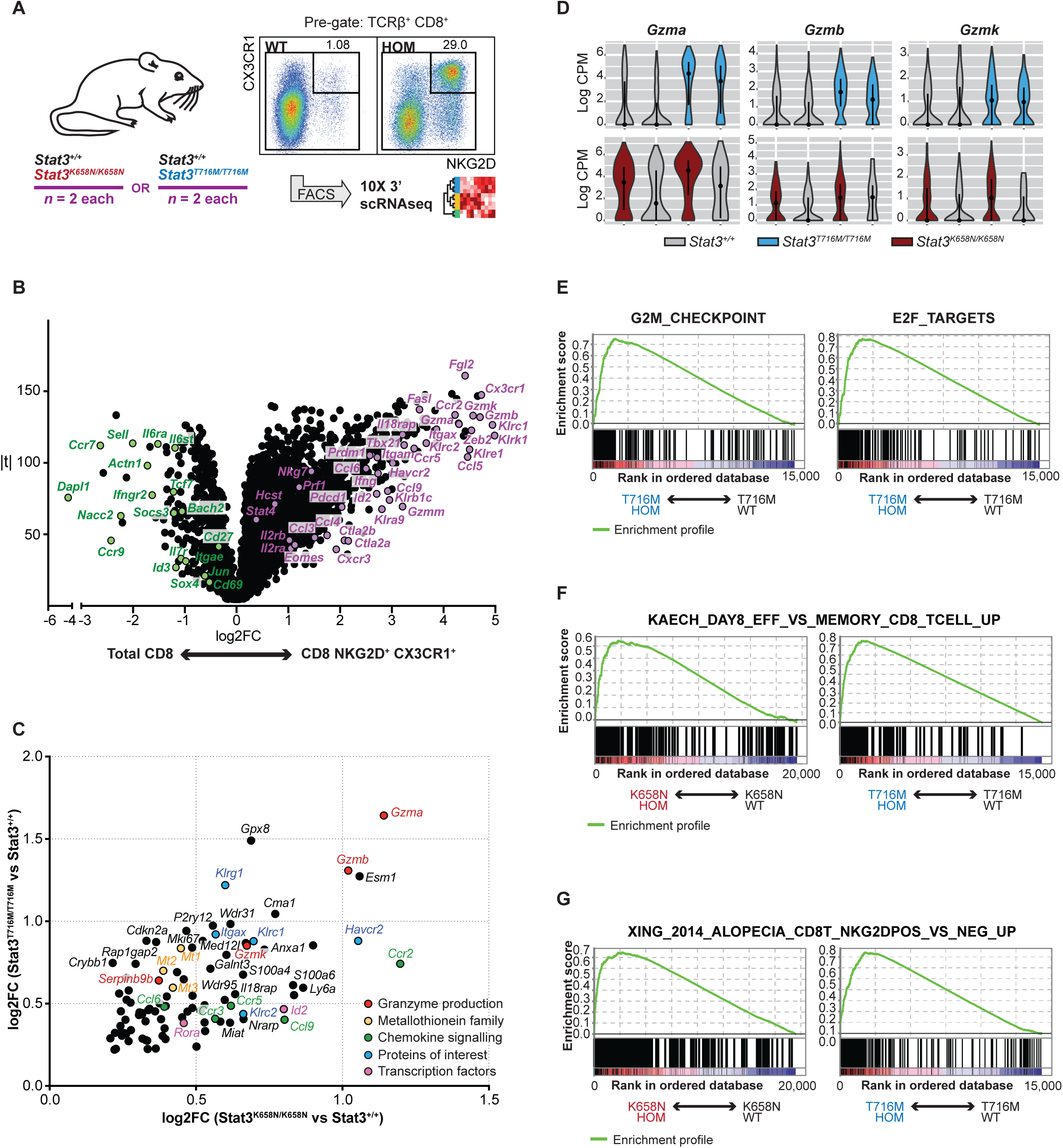
STAT3 GOF NKG2D^+^ CX3CR1^+^ CD8 T cells in mice overexpress a suite of killer cell genes. **A.** Schematic workflow for analysing single NKG2D^+^ CX3CR1^+^ CD8 T cells and total CD8 T cells (excluding NKG2D^+^ CX3CR1^+^ T cells) from *n* = 2 *Stat3^+/+^*or *Stat3^K658N/K658N^*and *n* = 2 *Stat3^+/+^* or *Stat3^T716M/T716M^*mice by global 3’ mRNA transcriptome scRNA-seq. **B.** Volcano plot of log2 expression fold-change (log2FC) versus moderated *t*-statistic for differentially expressed genes (family-wise error rate FWER < 0.05, using *limma*) in NKG2D^+^ CX3CR1^+^ CD8 T cells relative to total CD8 T cells in *Stat3^T716M/T716M^*mice. Text denotes genes of interest that were increased (purple) or decreased (green) in CD8 NKG2D^+^ CX3CR1^+^ T cells relative to total CD8 T cells. **C.** Dot plot of genes significantly up-regulated (FWER < 0.05; log2FC > 0.2) in both *Stat3^K658N/K658N^*(x-axis) and *Stat3^T716M/T716M^*(y-axis) NKG2D^+^ CX3CR1^+^ CD8 T cells relative to matched *Stat3^+/+^*NKG2D^+^ CX3CR1^+^ CD8 T cells. Coloured text denotes genes of interest. **D.** Violin plots showing kernel density estimations of *Gzma*, *Gzmb* and *Gzmk* mRNA log2 normalised counts per million (CPM) per cell in CD8 NKG2D^+^ CX3CR1^+^ T cells from mice of the indicated genotypes. Dots and lines indicate geometric median and interquartile range, respectively. **E-G.** Gene set enrichment analysis showing enrichment score (y axis) for rank-ordered genes (x axis) in *Stat3^K658N/K658N^*versus *Stat3^+/+^*and *Stat3^T716M/T716M^*versus *Stat3^+/+^*CD8 NKG2D^+^ CX3CR1^+^ T cells. **E.** HALLMARK gene sets G2M_CHECKPOINT (#M5901) or E2F_TARGETS (#M5925). **F.** Immunologic term gene set KAECH_DAY8_EFF_VS_MEMORY_CD8_TCELL_UP (#M3027). **G.** A published gene set of mRNAs increased in NKG2D^+^ Compared to NKG2D^-^ CD8 T cells from a mouse model of autoimmune alopecia (Xing et al., 2014).

The consequence of STAT3 GOF within this killer subset was explored by comparing *Stat3*-mutant and wild-type CD8 NKG2D^+^ CX3CR1^+^ T cells for differential gene expression. 1918 genes were significantly (family-wise error rate FWER < 0.05) increased and 571 decreased in *Stat3^T716M/T716M^* (**Supplementary Table 5**), and 216 genes increased and 18 decreased in *Stat3^K658N/K658N^*relative to *Stat3^+/+^*CD8 NKG2D^+^ CX3CR1^+^ T cells (**Supplementary Table 6**). Of these, 94 genes had mRNA increased by log2FC > 0.2 in both STAT3 GOF mutants (**Figure 5C**), including genes encoding inflammatory chemokines *Ccl6* and *Ccl9*, receptors for chemokines (*Ccr3, Ccr5, Ccr2*), metallothioneins (*Mt1, Mt2, Mt3*), transcription factors (*Id2, Rora*) and molecules crucial for effector or cytotoxic functions (*Gzma, Gzmb, Gzmk, Serpinb9b, Il18rap, Il18r1*; **Figure 5C,D**). It is notable that *Gzmb* has been shown to be a STAT3 target gene in activated CD8 T cells (Lu et al., 2019).

Gene set enrichment analysis (GSEA) comparing *Stat3*-GOF versus wild-type CD8 NKG2D^+^ CX3CR1^+^ T cells revealed a significant skew (FWER < 0.05) in both *Stat3^T716M/T716M^*and *Stat3^K658N/K658N^*mutant cells towards increased expression of HALLMARK gene sets associated with the G2M checkpoint and progression through the cell division cycle (#M5901) or with cell-cycle-related targets of E2F transcription factors (#M5925) (**Figure 5E**; **Supplementary Table 7**). *Stat3*-GOF cells also had significantly skewed increased expression of a gene set up-regulated during viral infection in effector relative to naïve CD8 T cells (#M3013 (Kaech et al., 2002)), including: *Acyp2, Adam8, Anxa1, Ccr2, Ccr5, Dock5, Errfi1, Galnt3, Gzma, Gzmb, Gzmk, Id2, Il18rap, Itgax, Kif11, Klrg1, Med12l, Mki67, Perp, Prc1, Rom1, Rora, Serpinb9b, S100a4, S100a6, Top2a* (**Supplementary Table 7**). *Stat3*-GOF cells also had significantly skewed increased expression of genes up-regulated in effector relative to memory CD8 T cells during peak viral infection (#M3027 (Kaech et al., 2002)) (**Figure 5F**, **Supplementary Table 7**).

Finally, we generated a custom immunological terms gene set comprising significantly increased mRNAs identified by Xing *et al*. in bulk RNA sequencing of CD3^+^ CD8^+^ CD44^+^ T cells that were NKG2D^+^ versus those that were NKG2D^-^, sorted from lymph nodes of autoimmune alopecic C3H/HeJ mice (Xing et al., 2014). Strikingly, this was the most significantly enriched of gene sets in *Stat3^K658N/K658N^*relative to *Stat3^+/+^*CD8 NKG2D^+^ CX3CR1^+^ T cells, and was also significantly enriched in *Stat3^T716M/T716M^*cells (**Figure 5G**, **Supplementary Table 7**).

We used flow cytometric analysis to validate selected gene expression changes at the protein level in single cells. Relative to wild-type counterparts, *Stat3^T716M/T716M^*CD8^+^ NKG2D^+^ CX3CR1^+^ T cells expressed significantly higher levels of *Cx3cr1* mRNA (**Supplementary Figure 4A**) and CX3CR1 cell-surface protein levels (**Supplementary Figure 4B**). Similarly, *Stat3*-mutant CD8 NKG2D^+^ CX3CR1^+^ T cells expressed significantly higher *Klrc1* (encodes NKG2A), *Klrc2* (encodes NKG2C) and *Klrg1* mRNA levels and correspondingly higher NKG2A/C/E and KLRG1 cell-surface levels (**Supplementary Figure 4C-E**). Thus, germline STAT3 GOF causes accumulation in both mice and humans of NKG2D^+^ effector CD8 T cells over-expressing mRNAs and proteins required for inflammatory effector and killer functions.

### Accumulating NKG2D^+^ CD8 T cells in STAT3 GOF mice comprise many expanded clones

Given that nearly half of human T-LGL’s have *STAT3* GOF somatic mutations (Jerez et al., 2012, Koskela et al., 2012, Fasan et al., 2013, Teramo et al., 2020), a key question was whether or not the accumulating NKG2D^+^ CD8 cells in *Stat3* GOF mice comprised a single leukemic clone. In wild-type and *Stat3*-GOF mice, the proportions of CD8 T cells expressing NKG2D were significantly higher in bone marrow relative to spleen (**Figure 6A**), and increased with age in spleen and blood (**Figure 6B**) but did not accumulate in overall frequencies consistent with leukemia. To analyse clonality, we performed TCR mRNA deep sequencing on splenic NKG2D^-^ and NKG2D^+^ CD8 T cells sorted by FACS from *Stat3^T716M^* mice (30-45 weeks old) and *Stat3^K658N^*mice (10-20 weeks old). NKG2D^+^ CD8 T cells from STAT3 GOF or wild-type mice were highly polyclonal, although the most frequent clonotype accounted for on average 26.40% (range 5.58 –88.2%) relative to 14.70% (range 2.06 – 47.70%) of unique *TCRA* reads in mutant relative to wild-type mice, respectively (**Figure 6C**). In mutant and wildtype mice, 50% of unique TCR reads came from less than 50 different clonotypes (**Figure 6C**) indicating that many large clones contribute to this subset. *Stat3*-mutant CD8 NKG2D^+^ T cells therefore did not constitute a clonal neoplasm, but the presence of many expanded clones among the CD8 NKG2D^+^ T cells was reflected by less diversity (as measured by Shannon entropy) compared to CD8+ NKG2D^-^ T cells (**Figure 6D**). A similar pattern of clonotype distribution frequencies was also observed following *TCRA* and *TCRB* deep sequencing of NKG2D^+^ CX3CR1^+^ CD8 T cells from *Stat3^K658N/K658N^*mice (**Figure 6E**). These results parallel the presence of many expanded clones in non-naïve CD8 T cell clusters 0, 7 and 11 in *STAT3* GOF patients (Figure 4), and resembles observations for CD57^+^ relative to CD57^-^ CD8 T cells in humans (Morley et al., 1995). Given that the total number of NKGD2^hi^or CD57^+^ CD8 cells was dramatically increased in mice or humans with germline *STAT3* GOF, these results indicate that these mutations are insufficient alone to cause monoclonal T-LGL but enhance accumulation of many large effector CD8 T cell clones.

**Figure 6.**
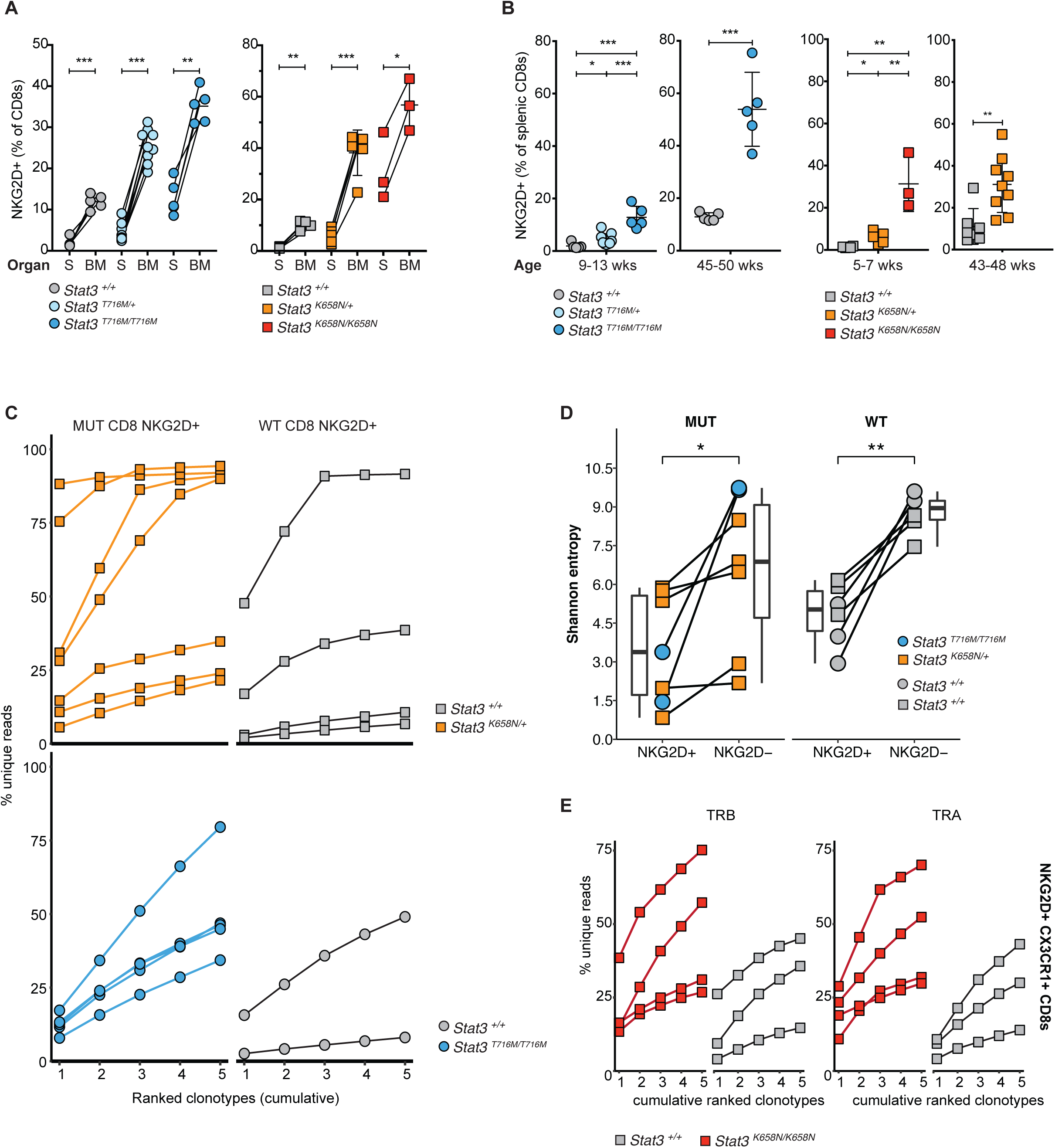
Oligoclonal *Stat3*-mutant CD8 NKG2D^+^ T cells accumulate in the spleen, blood and bone marrow with age. **A.** Percentage of CD8 T cells expressing NKG2D, in the spleen or bone marrow of individual mice with the indicated genotypes. Measurements made within same mouse are linked by a solid line. Statistical comparisons made by paired *t*-test. **B.** Percentage of splenic CD8 T cells expressing NKG2D, in 9-13 or 45-50 week old *Stat3^T716M^*, or 5-7 or 18-24 week old *Stat3^K658N^*mice of the indicated genotypes. Statistical comparisons made by *t*-test, corrected for multiple comparisons using the Holm-Sidak method. **C,D.** Following sorting of NKG2D^-^ and NKG2D^+^ splenic TCRβ^+^ CD8^+^ T cells, mRNA was prepared from cell pools and TCRα rearrangements were deep sequenced by 5’RACE from the *Tcra* constant region gene. **C.** Cumulative percentage of unique reads (y-axis, determined using unique molecular identifiers UMIs) captured by top 5 ranked clonotypes (x-axis), for cells from mice of the indicated genotypes. Lines join data points from individual mice. **D.** Shannon Entropy calculations of repertoire diversity, for cells from mice of the indicated genotypes. Lines join data points from the same mouse. Boxplots show median and interquartile range values. **E.** Cumulative percentage of unique reads (y-axis) captured by top 5 ranked clonotypes (x-axis), following sorting of NKG2D^+^ CX3CR1^+^ CD8 T cells from individual *Stat3^+/+^*(grey) and *Stat3^K658N/K658N^*(red) mice and 5’RACE *Tcrb* and *Tcra* deep-sequencing. *p* values are independent, calculated by Wilcoxon rank-sum test. * *p* < 0.05; ** *p* < 0.01; *** *p* < 0.001. A,B. Data are represented as mean ± SD.

### NKG2D and IL-2/IL-15 receptors drive accumulation of NKG2D^+^ CD8 T cells in STAT3 GOF mice

To identify which pathways are essential to drive the polyclonal expansion of STAT3 GOF CD8 NKG2D^+^ T cells, we tested whether their accumulation in *Stat3* GOF mice was diminished by genetically or pharmacologically interfering with CX3CR1, IL10R, NKG2D, IL6R, IL21R, IL7R or IL2/IL15R. Homozygous germline *Cx3cr1* deletion resulted in a variable, small decrease in percentage of splenic NKG2D^+^ CD8 T cells in *Stat3^K658N^*mutant mice (**Supplementary Figure 5A**). CX3CR1 expression on CD8 NKG2D^+^ T cells is largely restricted to a subpopulation with a CD27^low^KLRG1^high^CD62L^neg^(and CD127^low^CD44^high^CXCR3^low^) profile (**Supplementary Figure 1D**, **Supplementary Figure 5B,C**), but we observed no change in the percentage of this subset between *Cx3cr1^+/+^*, *Cx3cr1^KO/+^*and *Cx3cr1^KO/KO^ Stat3*-mutant mice (**Supplementary Figure 5D**). CX3CR1 therefore appears dispensable for the accumulation of most NKG2D^+^ CD8 T cells with overactive STAT3 in blood, spleen and bone marrow.

The STAT3-signalling cytokine IL-10 decreases inflammation and antigen-induced CD8 T cell activation (Smith et al., 2018). However, in the absence of the essential alpha chain of the receptor for IL-10, in *Stat3^T716M/T716M^Il10ra^KO/KO^*relative to *Stat3^T716M/T716M^ Il10ra^+/+^*mice, there was no significant change in the mean percentage of NKG2D^+^ Cells within CD8+ effector memory T cells (Supplementary Figure 5E).

To determine if NKG2D signalling drives the accumulation of *Stat3*-mutant effector CD8 T cells, we crossed *Stat3*^Δ^*^K658N^*mice to *Klrk1* mice lacking NKG2D. As a proxy marker for NKG2D-expressing T cells, we took advantage of the fact that most NKG2D^+^ CD8 T cells in *Stat3* GOF mice co-express NKG2A/C/E (**Supplementary Figure 5F**). *Klrk1^KO/KO^ Stat3^K658N/K658N^*mice had half as many NKG2A/C/E^+^ CX3CR1^+^ Cells as a percentage of CD8 T cells or of total leukocytes in spleen, bone marrow and blood compared with *Stat3^K658N/K658N^*mice with intact NKG2D (**Figure 7A** and data not shown). NKG2D therefore contributes to *STAT3*-mutant effector CD8 T cell accumulation, but NKG2D deficiency is insufficient to fully suppress accumulation of these cells in *Stat3* GOF compared to *Stat3* wild-type mice.

**Figure 7.**
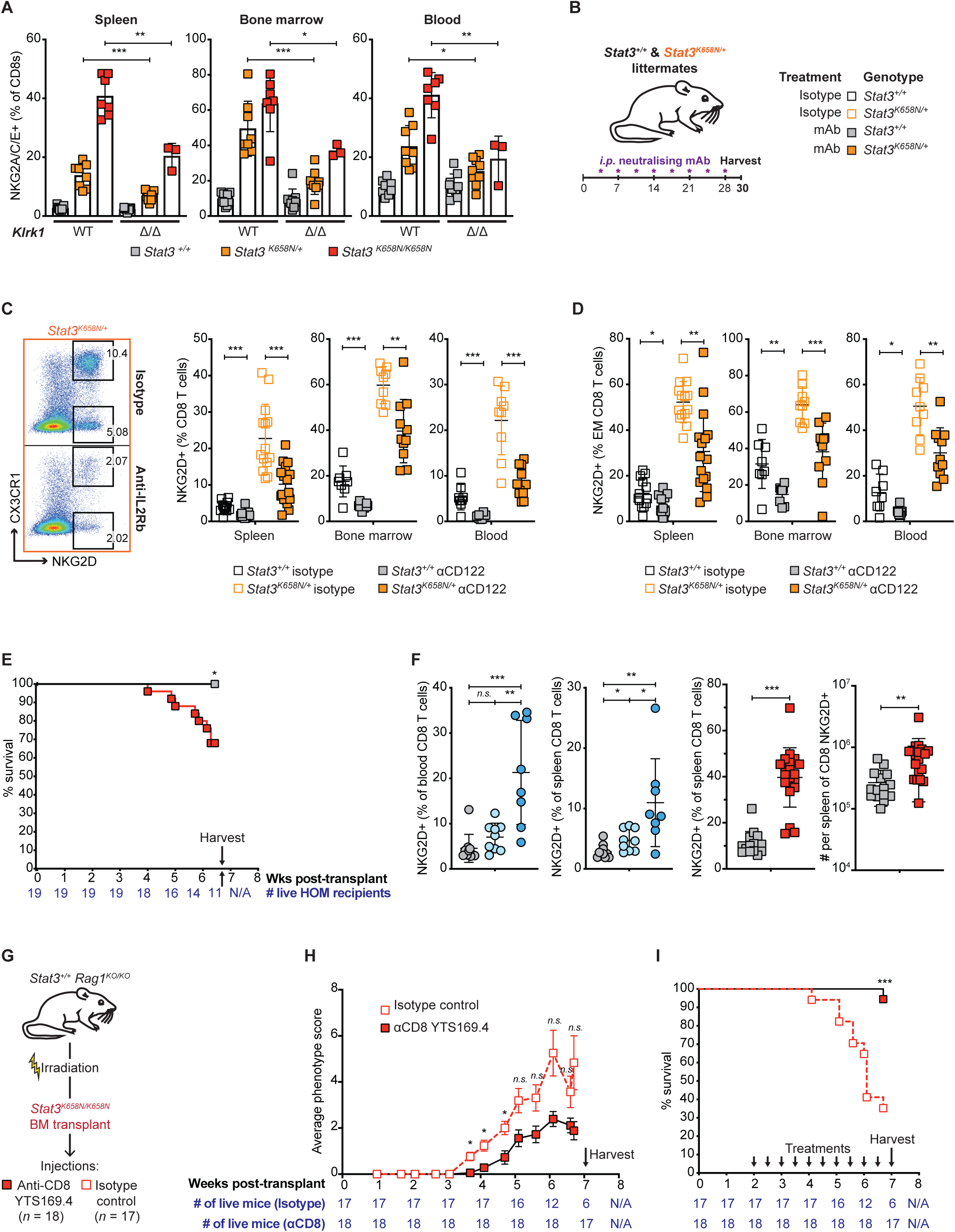
Essential roles of NKG2D and IL2RB in STAT3 GOF effector CD8 cell accumulation and of CD8 T cells in lethal pathology. **A.** Percentage of NKG2A/C/E^+^ Cells among CD8 T cells in the spleen of individual *Stat3^+/+^*, *Stat3^K658N/+^*or *Stat3^K658N/K658N^*mice that were *Klrk1^+/+^*(WT) or *Klrk1^KO/KO^*(Δ/Δ). **B.** Experimental design for treating *Stat3^+/+^*or *Stat3^K658N/+^*mice with 8 bi-weekly intra-peritoneal (*i.p.*) injections of blocking monoclonal antibody (mAb) to different receptors or control antibody (isotype). **C.** Representative flow cytometry plots of spleen CD8 T cells in *Stat3^K658N/+^*mice showing percentage expressing NKG2D, and values for individual *Stat3^+/+^*or *Stat3^K658N/+^*mice treated with isotype or blocking anti-CD122 mAb. **D.** As in C, but showing percentage NKG2D^+^ among CD44^+^ CD62L^-^ effector memory CD8 T cells. **E.** Kaplan-Meier survival curve for recipients of *Stat3^+/+^*(black, n=15) or *Stat3^K658N/K658N^*(red, n= 19) marrow. **F.** Frequency of NKG2D^+^ Cells among CD8 T cells in the blood or spleen of individual recipients of *Stat3^+/+^*(grey), *Stat3^T716M/+^*(light blue), *Stat3^T716M/T716M^*(dark blue), *Stat3^K658N/+^*(orange) or *Stat3^K658N/K658N^*(red) marrow. **G.** Experimental design for selectively depleting STAT3 GOF CD8 T cells from recipients of *Stat3^K658N/K658N^*marrow with bi-weekly injections of anti-CD8 or isotype control antibodies beginning 2 weeks post-transplant. **H.** Mean phenotype score of the mice in (F). Scores for each recipient were calculated as equal weighted averages of severity scores on a scale from 0-3 (0 = undetectable; 1 = minor; 2 = moderate; 3 = major) for: (1) hair loss or inflammation around eyes, or cataracts; (2) skin inflammation or ‘flakiness’ on snout or face; (3) skin inflammation or ‘flakiness’ on ears; (4) skin inflammation or hair loss, primarily ventral, progressing to alopecia; (5) skin inflammation or flakiness on tail, or ringtail. **I.** Kaplan-Meier survival curve for *Stat3^K658N/K658N^*bone marrow recipients 0-7 weeks post-transplant (Tx) treated with anti-CD8 (black line, red fill) versus isotype control (dashed line, no fill). Numbers of mice in both treatment arms indicated below. **A.** Data pooled from *n* = 2 independent experiments. **C,D.** Data pooled from *n* = 3 independent experiments. Statistical comparisons made by *t*-test, corrected for multiple comparisons using the Holm-Sidak method. **E,I.** Statistical analysis calculated using the product-limit method on GraphPad Prism, accounting for censored mice. Statistical significance calculated by log-rank test. **F.** Statistical comparisons in by *t*-test, corrected for multiple comparisons using the Holm-Sidak method. Data representative of *n* > 2 independent experiments with *n* > 5 mice per group. **G**. Statistical analysis calculated by 2-way ANOVA followed by Šidák’s multiple comparisons test. * *p* < 0.05; ** *p* < 0.01; *** *p* < 0.001. A,C,D,F. Data are represented as mean ± SD.

Pharmacologic interference experiments were performed by treating *Stat3^K658N^* heterozygous mice with monoclonal antibodies (mAbs) known to block the function of different cell-surface receptors (**Figure 7B**). Consistent with the results of genetic interference of NKG2D in *Klrk1^KO^*mice, *Stat3^K658N^*mice treated with a blocking anti-NKG2D mAb had a mean 42% fewer NKG2A/C/E^+^ CX3CR1^+^ CD8 T cells than mice treated with a control mAb (**Supplementary Figure 5G**). Blocking IL-6 or IL-21 signalling to STAT3 or blocking IL-7 signalling to STAT5, by treating with anti-IL6R, anti-IL21R or anti-IL7R mAbs, had no significant effect on the mean % NKG2D^+^ CD8 T cells in the spleen, bone marrow and blood of *Stat3*-mutant mice compared to treatment with control mAbs (**Supplementary Figure 5H, I**). By contrast, *Stat3^K658N^*mice treated with anti-CD122 blocking mAb, targeting the IL-2Rβ chain required for IL-15 and IL-2 signalling, had half as many NKG2D^+^ Cells among all CD8 T cells (**Figure 7C**) or among CD62L^-^ CD44^+^ effector memory CD8 T cells (**Figure 7D**) in the spleen, bone marrow and blood compared to control mAb-treated animals. Collectively, these findings establish that NKG2D and IL-15/IL-2 signals are required but IL-10, IL-6, IL-21 or IL-7 signals appear individually dispensable for the exaggerated accumulation of NKG2D^+^ CD8 T cells harbouring pathogenic *STAT3* GOF mutations.

### Depleting *STAT3* GOF CD8 T cells ameliorates inflammation and lethality

Pathology in humans and mice with germline *STAT3* GOF syndrome may be due to aberrant STAT3 signalling and function in non-hematopoietic cells. We therefore generated chimeric mice with hematopoietic-restricted *Stat3* GOF mutations by transplanting *Stat3^+/+^ Rag1^KO/KO^*mice with *Stat3*-GOF or wild-type bone marrow (**Supplementary Figure 6A**). Mice transplanted with *Stat3*-GOF but not wild-type marrow developed a range of gross pathologies (**Supplementary Figure 6B**): recipients of *Stat3^T716M/T716M^*marrow primarily developed lymphadenopathy, splenomegaly, and/or cataracts; recipients of *Stat3^K658N/K658N^*marrow developed lymphadenopathy, splenomegaly, hair loss around the eyes, cataracts, ventral hair loss, diarrhea, weight loss (wasting) requiring ethical culling, or were unexpectedly found dead. 8 of 25 recipients of *Stat3^K658N/K658N^*marrow required sacrifice at ethical end-point within 7 weeks post-transplant compared to 0 of 15 recipients of *Stat3^+/+^*marrow (**Figure 7E**). NKG2D^+^ CD8 T cells were increased in recipients of *Stat3^T716M/T716M^* or *Stat3^K658N/K658N^* bone marrow (**Figure 7F**).

To test the role of the CD8^+^ T cells bearing *STAT3* GOF in the pathology that developed in *Stat3^K658N/K658N^*marrow transplant recipients, CD8 T cells were selectively depleted by injections of a depleting anti-CD8 mAb or isotype control mAb starting 2 weeks after bone marrow transplant (**Figure 7G**). *Stat3^K658N/K658N^*marrow recipients in the control arm started developing gross pathology 3-5 weeks post-transplant at the time that mature T cells reconstitute from hematopoietic stem cells, but the pathology was significantly diminished in recipients treated with anti-CD8 (**Figure 7H**). By 7 weeks post-bone marrow transplantation, 11 of 17 recipients in the isotype control arm had to be ethically culled for weight loss and wasting disease compared to only 1 out of 18 recipients in the CD8 depleted arm (**Figure 7I**). Thus, STAT3 GOF limited to blood cells is sufficient to cause accumulation of NKG2D^+^ CD8 T cells and lethal pathology, and CD8 cells are required for this pathology.

## DISCUSSION

The findings here illuminate a pathogenic mechanism in autoimmune disease by resolving a key question about the intersection between cancer and autoimmunity: whether the *STAT3* GOF somatic mutations often found in CD8 T cells in T-LGL are a cause or an effect of clonal expansion and accompanying autoimmune disease. The presence of circulating clones of CD8 T cells bearing *STAT3* GOF somatic mutations is strongly correlated with rheumatoid arthritis in T-LGL (Koskela 2012; Rajala 2015), with Felty syndrome of rheumatoid arthritis, neutropenia and splenomegaly (Savola et al., 2018), and with primary red cell aplasia (Kawakami et al., 2018). However, the size of these STAT3 mutant clones varies widely, with a mutated STAT3 variant allele frequency (VAF) in CD8 cells ranging from 0.6-51% and median 17% in T-LGL (Rajala 2015), 1-8% in Felty syndrome (Savola et al., 2018), and 1.1-16% and median 2.5% in primary red cell aplasia (Kawakami et al., 2018). Clones representing 20% of CD8 cells (10% VAF for heterozygous mutations) are within the physiological range for effector memory CD8 T cells (Posnett et al., 1994) and not in the magnitude of malignant leukemias. Moreover, *STAT3* GOF mutations are found in healthy people with VAFs of 0.4-1.9% in CD8 cells, and in asymptomatic HTLV-infected people with VAFs of 0.5-11.9% in CD8 cells (Valori et al., 2021, Kim et al., 2021). Here we show that STAT3 GOF mutations in the germline of humans or mice do not perturb the preponderant population of naïve CD8 T cells, consistent with previous experiments by retroviral transduction of mouse bone marrow (Couronne et al., 2013, Dutta et al., 2018), but instead they selectively drive indolent expansion of a normally minor subset of activated effector CD8 T cells closely resembling T-LGL cells. Like T-LGL, these *STAT3* GOF T cells strongly expressed the genes for cytotoxic granule proteins (granzymes, perforin) and there was high variability in the size of the expanded clones and accompanying autoimmune pathology. The results show that CD8 T cells bearing STAT3 GOF mutations cause lethal inflammatory disease. By revealing that these cells express high levels of NKG2D and DAP10*/HSCT* and depend upon NKG2D and the IL-15 receptor IL2Rβ for their accumulation, the findings here bring together diverse evidence explaining their indolent clonal expansion, NK-like effector gene expression profile, and connection to autoimmune disease (**Graphical Abstract**) as discussed below.

The diminished accumulation of STAT3 GOF CD8 effector T cells when NKG2D was genetically eliminated or blocked by antibody in the experiments here is consistent with the evidence that NKG2D is a costimulatory receptor that activates PI3-kinase signalling in T cells similarly to CD28 and ICOS, albeit through a YXXM motif in the heterodimeric partner protein DAP10 (*HCST*; (Wu et al., 1999)). Combined triggering by TCR and NKG2D enhances human and mouse CD8 T cell proliferation in vitro (Groh et al., 2001, Jamieson et al., 2002) and increases clonal expansion of autoimmune CD8 T cells bearing TCRs against an MHC I-presented peptide fragment of pancreatic islet-specific glucose-6-phosphatase catalytic subunit related protein (IGRP) in NOD mice (Ogasawara et al., 2004).

IL2Rβ blockade also diminished accumulation of STAT3 GOF CD8 effector T cells in our experiments, and this result is consistent with the role of IL-15 in CD8 effector T cell accumulation. IL-15 is presented to T cells as a heterodimer with IL-15Rα on the surface of IL-15-expressing cells to engage IL-2Rβ and IL-2Rγ on T cells and activate STAT3 and STAT5 (Waldmann et al., 2020). Exogenously-administered IL-15/IL-15RA heterodimers are potent drivers of homeostatic CD8 T cell proliferation in vivo (Chertova et al., 2013) and IL-15 and IL-15RA expression by dendritic cells and monocytes is essential for the accumulation of circulating effector memory CD8 T cells (Ku et al., 2000, Zhang et al., 1998, Stonier et al., 2008). Large monoclonal or oligoclonal CD8 T cell expansions develop spontaneously in old mice and their indolent cell division is inhibited by antibody blockade of IL2Rβ (Ku et al., 2001), although it is not known if these clones have acquired *STAT3* GOF somatic mutations.

Within TCR-activated CD8 T cells, IL-15 induces expression of NKG2D and *HSCT/*DAP10 (Roberts et al., 2001, Meresse et al., 2004), as does high concentrations of IL-2 which engages IL2Rβ and IL2Rγ with lower affinity than IL-15/IL-15RA heterodimers (Verneris et al., 2004, Jabri and Abadie, 2015). NKG2D induction is likely mediated by STAT3, given that the *KLRK1* gene is a direct STAT3 target in NK cells (Zhu et al., 2014) and that NKG2D was more highly expressed on CD8 T cells from STAT3 GOF patients (**Figure 3D**). Interferon gamma, which was a highly expressed mRNA in the NKG2D^hi^STAT3 GOF effector CD8 T cells analysed here, induces IL-15 expression in macrophages (Doherty et al., 1996) and hair follicles (Xing et al., 2014). Thus, *STAT3* GOF mutations may exaggerate a physiological positive feedback loop where IL-15 promotes accumulation of NKG2D^hi^IFNγ-expressing effector CD8 T cells which in turn induce IL-15 (**Graphical Abstract**). This loop may be further exaggerated by another STAT3 target gene, *CCL5* (Yang et al., 2007), found highly expressed in STAT3 GOF CD8 T cells. *CCL5* encodes the T cell-attracting chemokine RANTES and is induced by NKG2D stimulation of TCR-activated CD8 T cells to recruit other T cells (Markiewicz et al., 2012).

How does the exaggerated accumulation of *STAT3* GOF NKG2D^hi^granzyme^+^ perforin^+^ effector CD8 T cells contribute to autoimmune disease? This likely reflects an interplay between the induction of NKG2D ligands and IL-15 by cellular stress in the autoimmune targeted organs (Jabri and Abadie, 2015) and NKG2D-stimulated T cells having a lower threshold for TCR-mediated IFNγ production and cytolysis (Roberts et al., 2001, Billadeau et al., 2003) or displaying MHC-independent NK-cell like cytotoxicity (lymphokine activated killer cells/ bystander killing (Verneris et al., 2004, Karimi et al., 2005, Meresse et al., 2004). The costimulatory ligands for NKG2D comprise two families of cell surface proteins evolutionarily related to MHC Class I (Bauer et al., 1999, Wu et al., 1999, Raulet et al., 2013). The first family comprises MHC class I chain-related A and B proteins (MICA, MICB) and is absent in rodents. The second comprises products of six human and nine mouse genes encoding Retinoic Acid Early Transcripts (RAETs), UL16-binding proteins (ULBPs), murine UL16-binding protein like transcript (MULTI), and Histocompatibility 60 proteins (H60). Expression of genes encoding these NKG2D ligands is induced on cells that are infected, have undergone DNA damage, have oncogenic mutations, or have been exposed to inflammatory cytokines (Raulet et al., 2013).

The high frequency of rheumatoid arthritis in T-LGL patients with *STAT3* GOF mutant CD8 T cell clones (Koskela et al., 2012, Rajala et al., 2015) is likely connected to the high expression of MICA and other NKG2D ligands on inflamed synovium in rheumatoid arthritis (Groh et al., 2003). NKG2D expression on T cells is normally controlled by ligand-induced down-modulation but this is reversed by IL-15 (Groh et al., 2002, Roberts et al., 2001) which also induces NKG2D ligands on synoviocytes (Groh et al., 2003). IL-15 levels are increased in the synovial fluid and membrane of patients with rheumatoid arthritis (McInnes et al., 1996) and in the serum of patients with T-LGL (Chen et al., 2012).

With respect to the dermatitis and acanthotic keratinocyte hyperproliferation caused by *STAT3* GOF CD8 T cells here, it is relevant that the NKG2D-ligand H60c is selectively expressed in skin keratinocytes and induced by wounding to activate NKG2D-PI3K signalling, cytotoxic granule release and cytokine production by intraepithelial T cells (Whang et al., 2009, Ibusuki et al., 2014). In celiac disease, MICA expression is highly induced on intestinal epithelial cells by dietary gluten through IL-15. Epithelial MICA returns almost to baseline on a gluten-free diet, except in patients with refractory celiac disease type 2 who have developed clonal expansions of innate lymphocytes that often carry *STAT3* GOF mutations (Meresse et al., 2004, Ettersperger et al., 2016, Cording et al., 2021, Soderquist et al., 2021, Hue et al., 2004).

People with germline *STAT3* GOF mutations can develop neonatal type 1 diabetes (Flanagan et al., 2014). It has recently been shown that one of these *STAT3* GOF mutations (K392R in the DNA binding domain), when introduced into the germline of diabetes-prone NOD mice, accelerates autoimmune diabetes by increasing the frequency of CD8^+^ CD44^hi^CD62L^-^ effector T cells expressing granzymes and *Ccl5*/RANTES (Warshauer et al., 2021). NKG2D expression or function was not examined, but autoimmune *Stat3^K392R^*CD8 T cell dysregulation likely also involves NKG2D because the IGRP-reactive CD8 T cells that mediated accelerated diabetes have previously been shown in NOD mice to depend upon NKG2D for their accumulation in and destruction of pancreatic islets (Ogasawara et al., 2004). In NOD mice, *Raet1* transcripts are constitutively expressed by pancreatic islet beta cells independent of any T or B cell autoimmune reaction (Ogasawara et al., 2004), possibly reflecting the polygenic inheritance of unfolded protein stress in the endoplasmic reticulum of islet beta cells in this strain (Dooley et al., 2016). Transgenic RAE1 expression restricted to pancreatic islet cells is sufficient to recruit killer CD8 T cells to the islets in a DAP10-dependent and TCR-independent manner (Markiewicz et al., 2012).

Collectively, the findings here support a model where autoimmunity associated with *STAT3* GOF somatic mutations in CD8 T cells reflects exaggeration by these mutations of a feedforward loop that has three key drivers (**Graphical Abstract**): (1) target cells that have upregulated NKG2D ligands due to inherited, somatically acquired, or environmentally induced abnormalities; (2) target tissues and adjacent macrophages and dendritic cells that have upregulated IL-15 in response to these abnormalities and to effector CD8 T cell infiltration and activation; and (3) upregulation of NKG2D and its signalling partner DAP10/*HCST* in the infiltrating CD8 T cells. This loop provides many points where inherited or environmental factors can tip clonal expansion of NKG2D^hi^effector CD8 T cells into pathological destruction of neutrophils, erythrocytes, joint synovium, islet beta cells, or epithelial barriers in the skin and intestine. The results here highlight the need for deeper testing for CD8 T cell clones with *STAT*3 GOF somatic mutations in autoimmune disease, and stratified clinical trials of inhibitors of the STAT3/NKG2D/IL15 loop in T-LGL and autoimmune disease patients where these rogue CD8 clones are present.

## ACKNOWLEDGEMENTS

We thank the patients and their families. We thank the Garvan-Weizmann Centre for Clinical Genomics (Garvan Institute of Medical Research, Sydney, Australia), the Garvan Biological Testing Facility and the Phenomics Australia Histopathology and Slide Scanning Service, for providing technical services. This work was supported by National Health and Medical Research Council (NHMRC) Program (1113904, to C.C.G.) and Fellowship (1081858, to C.C.G.) grants and by The Bill and Patricia Ritchie Foundation.

## AUTHOR CONTRIBUTIONS

E.M-F designed and performed the majority of experiments; R.B. generated CRISPR/Cas9 *Stat3*-mutant mouse models; K.P. and G.R. performed flow cytometry on human PBMCs; D.S, G.U, I.C, J.W.L, K.H, K.P, L.K, M.O, M.A.C, M.R.J.S, S.M, S.B, S.G.T obtained patient peripheral blood samples; M.S. and G.A. aided in generating TCR deep sequencing or single-cell multi-omics data; G.A., K.J.L.J. and T.J.P performed bioinformatics analyses; E.M-F, S.G.T, J.H.R and C.C.G interpreted experiments and wrote the manuscript.

## STAR METHODS

### RESOURCE AVAILABILITY

#### Lead contact

Further information and requests for resources and reagents should be directed to and will be fulfilled by the Lead Contact, Christopher C. Goodnow.

#### Materials availability

This study did not generate new unique reagents.

#### Data and code availability

• Single-cell RNA-seq and TCR mRNA deep sequencing data have been deposited in NCBI BioProject PRJNA804580. The accession number for these data is NCBI BioProject: PRJNA804580, as listed in the Key Resources table.

• Bioinformatic workflows used to analyse the data in this study are available upon request.

• Any additional information required to reanalyze the data reported in this paper is available from the lead contact upon request.

### EXPERIMENTAL MODEL AND SUBJECT DETAILS

#### Human subjects

This study was approved by the respective ethics review boards of the participating institutes, including the Ethics committees of Sydney Local Health District RPAH Zone Human Research Ethics Committee and Research Governance Office, Royal Prince Alfred Hospital, Camperdown, NSW, Australia (Protocol X16-0210/LNR/16/RPAH/257); the South East Sydney Local Health District Human Research Ethics Committee, Prince of Wales/Sydney Children’s Hospital, Randwick, NSW, Australia (Protocol HREC/11/POWH/152); the Helsinki University Hospital Ethics committee. Written informed consent for genetic investigations and immunological analyses, as well as publication of data, was obtained from each family.

#### Mouse handling, housing and husbandry

All mouse handling and experimental methods were performed in accordance with approved protocols of the Garvan Institute of Medical Research/St Vincent’s Hospital Animal Ethics Committee. All mice were bred and maintained in specific pathogen-free conditions at Australian BioResources (ABR; Moss Vale, Australia) or at the Garvan Institute of Medical Research Biological Testing Facility (BTF). Within independent experiments, *Stat3* wild-type and mutant animals were sex- and age-matched. All experiments conformed to the current guidelines from the Australian Code of Practice for the Care and Use of Animals for Scientific Purposes. Mice were genotyped by the Garvan Molecular Genetics (GMG) facility at the Garvan Institute of Medical Research.

#### Mouse anatomy and histopathology

Detailed anatomical and histopathology analysis was performed as a commercial service, on over 40 organs from *Stat3^T716M^*wild-type (*n*=3), homozygous (*n*=3), *Stat3^K658N^*wild-type (*n*=5), heterozygous (*n*=1) and homozygous (*n*=5) mice, by the Phenomics Australia Histopathology and Slide Scanning Service (University of Melbourne, Australia).

#### Mouse strains

*Stat3^T716M^*, *Stat3^K658N^, Il10ra^KO^*, *Klrk1^KO^*and *Cx3cr1^KO^*mice were produced by CRISPR/*Cas9* gene targeting in mouse embryos, following established molecular and animal husbandry techniques (Yang et al., 2014). Target gene-specific single guide RNAs (sgRNA; 15ng/μl) were microinjected into the nucleus and cytoplasm of mouse zygotes, together with polyadenylated S. pyogenes *Cas9* mRNA (30ng/μl) and a gene-specific 150 base single-stranded, deoxy-oligonucleotide homologous recombination substrate (15ng/μl). Injections were performed into C57BL/6 zygotes for each of these strains but the *Stat3^K658N^*strain, where injections were performed into C57BL/6J x FVB/N F1 zygotes. Founder mice heterozygous for alleles successfully modified by homologous recombination were back-crossed with syngeneic partners and then inter-crossed to establish the *Stat3^T716M^*or *Stat3^K658N^* mouse lines.

For *Stat3^T716M^*, the sgRNA was produced based on a target site in exon 23 (CAGGTCAATGGTATTGCTGC*<u>AGG</u>* = T716M, PAM italicised and underlined) of *Stat3*. The deoxy-oligonucleotide encoded the T716M (A<u>C</u>G><u>A</u>TG) substitution and a PAM-inactivating silent mutation in the T717 codon (AC<u>C</u>>AC<u>A</u>). For *Stat3^K658N^*, the sgRNA was produced based on a target site in exon 21 (CATGGATGCGACCAACATCC*<u>TGG</u>* = K658N, PAM italicised and underlined) of *Stat3*. The deoxy-oligonucleotide encoded the K658N (AA<u>G</u>><u>A</u>AC) substitution and a PAM-inactivating silent mutation in the L666 codon (CT<u>G</u>>C<u>T</u>C).

For *Il10ra^KO^*, the sgRNA was produced based on a target site exon 3 (GGTGAACGTTGTGAGATCAC*<u>AGG</u>*, PAM italicised and underlined) of *Il10ra*. The deoxy-oligonucleotide encoded the T86I (A<u>T</u>A>A<u>C</u>A) substitution and a PAM-inactivating silent mutation in the S82 codon (TC<u>A</u>>TC<u>C</u>). A founder carrying an 8bp frame shift mutation after the second base of the S82 codon was bred to establish the *Il10ra^KO^*line. For *Cx3cr1^KO^*, the sgRNA was produced based on a target site in the single coding exon, exon 2 (TACGCCCTCGTCTTCACGTT*<u>CGG</u>*, PAM italicised and underlined) of *Cx3cr1*. A founder mouse carrying an 1bp deletion that caused a frameshift after the T44 codon and an in-frame TGA stop signal 27 codons downstream was back-crossed with syngeneic partners to establish the *Cx3cr1^KO^*line. For *Klrk1^KO^*, the sgRNAs were produced based on target sites in intron 4 and the 3’ UTR (TCACAACGTGGTATAGTCCT*<u>AGG</u>* and GTTGAAGCCTATCCAAACTA*<u>GGG</u>*, PAM italicised and underlined) of *Klrk1*. A founder mouse that carried a 4,781 bp deletion in *Klrk1* that removed exons 4 to 8 (encoding the transmembrane and extracellular domains of NKG2D) was back-crossed with syngeneic partners and then inter-crossed to establish the *Klrk1^KO^*line. C57BL/6 JAusb (C57BL/6J), C57BL/6 NCrl, B6.SJL-*Ptprc^a^Pepc^b^*(CD45.1) and B6.129S7-*Rag1^tm1Mom^*/J (*Rag1^KO/KO^*) mice were purchased from ABR.

#### Chimeras

Age- and sex-matched *Stat3^+/+^ Rag1^KO/KO^*C57BL/6J mice were irradiated with one dose of 425 Rad from an X-ray source (X-RAD 320 Biological Irradiator, PXI) and injected intravenously with 2-6 x 10^6^ lineage-depleted donor bone marrow from *Stat3^T716M^*or *Stat3^K658N^*wild-type or homozygous mutant mice. The donor bone marrow was depleted of lineage-positive cells by MACS using a cocktail of antibodies to B220, CD3, CD4, CD8, CD11b, CD11c, CD19, LY-6C, LY-6G, NK1.1, TCRβ prior to injection. Recipients were sacrificed 8-14 weeks after transplantation to allow immune reconstitution.

#### *In vivo* monoclonal neutralising antibody treatments

Recipient *Stat3^K658N^*mice were mildly anesthetised by isoflurane and injected *i.p.* twice per week, to a total of 7 or 8 injections, with varying amounts of monoclonal antibodies as tabulated below, and mice analysed by flow cytometry 2-3 days after the last injection.

#### BioXCell monoclonal neutralising antibodies

**BioXCell monoclonal neutralising antibodies.**
| Cat number | Antibody | Clone | µg per injection |
| --- | --- | --- | --- |
| BE0111 | Armenian Hamster anti-mouse NKG2D | HMG2D | 200 |
| BE0091 | Polyclonal Armenian Hamster IgG | N/A | * |
| BE0298 | Mouse anti-mouse CD122 (IL-2Rβ) | TM-β1 | 200 |
| BE0047 | Rat IgG2b anti-mouse IL-6R | 15A7 | 250 |
| BE0090 | Rat IgG2b isotype control, anti-keyhole limpet hemocyanin | LTF-2 | * |
| BE0258 | Rat IgG2a anti-mouse IL-21R | 4A9 | 200 |
| BE0065 | Rat IgG2a anti-mouse IL-7Rα | A7R34 | 200 |
| BE0089 | Rat IgG2a isotype control, anti-trinitrophenol | 2A3 | * |

\* Isotype control antibodies were injected at equal concentration and final amount to the relevant treatment antibody.

#### Flow cytometry and cell-sorting

Peripheral blood mononuclear cells (PBMCs) were prepared from total blood collected from individuals with STAT3 GOF syndrome. In mice, single-cell suspensions were prepared from mouse spleen, bone marrow, inguinal lymph nodes, peritoneal cavity and blood. 1-4 x 10^6^ cells in PBS 2% FCS were transferred into appropriate wells of a 96-well U bottom plate. To prevent non-specific antibody binding, cells were incubated with F_c_ blocking antibody for 20 min at 4°C in the dark. Cells were then incubated with antibodies for 30 min, on ice and in the dark. To fix cells, they were incubated in 10% formalin (Sigma-Aldrich) for 15 min at 4°C, and washed and resuspended in PBS 2% FCS. To stain for intracellular nuclear proteins, cells were fixed and permeabilised using the manufacturer’s instructions and the eBioscience Transcription Factor Staining kit. Stained single-cell suspensions were acquired on the BD LSRFortessa^TM^.

Where appropriate, following extracellular antibody staining, immune populations were sorted by fluorescence-activated cell sorting (FACS) on a FACS Aria III (BD Biosciences).

#### Anti-mouse antibodies used for flow cytometric analyses

**Anti-mouse antibodies used for flow cytometric analyses.**
| Cat number | Antibody/dye | Fluorochrome | Company | Clone |
| --- | --- | --- | --- | --- |
| A1310 | 7AAD | N/A | Invitrogen | N/A |
| 100204 | CD3 | FITC | BioLegend | 17A2 |
| 100217 | CD3 | PerCP Cy5.5 | BioLegend | 17A2 |
| 563331 | CD4 | BV786 | BD Biosciences | GK1.5 |
| 564306 | CD4 | BUV737 | BD Biosciences | SK3 |
| 17-0051-81 | CD5 | APC | Thermo Fischer | 53-7.3 |
| 563796 | CD8 | BUV395 | BD Biosciences | 53-6.7 |
| 557654 | CD8 | APC Cy7 | BD Biosciences | 53-6.7 |
| 557396 | CD11b | FITC | BioLegend | M1/70 |
| 101208 | CD11b | PE | BioLegend | M1/70 |
| 45-0114-82 | CD11c | PerCP/Cy5.5 | eBioscience | N418 |
| 115546 | CD19 | BV510 | BioLegend | 6D5 |
| 115539 | CD19 | BV605 | BioLegend | 6D5 |
| 101820 | CD24 | Pacific Blue | BD Biosciences | M1/69 |
| 102030 | CD25 | PerCP Cy5.5 | BioLegend | PC61 |
| 558642 |  | PE | BD Biosciences | 7D4 |
| 557192 |  | APC | BD Biosciences | PC61 |
| 124216 | CD27 | PE Cy7 | BioLegend | LG.3A10 |
| 122007 | CD28 | FITC | BioLegend | E18 |
| 562768 | CD38 | BV421 | BD Biosciences | Ab90 |
| 563058 | CD44 | BV605 | BD Biosciences | IM7 |
| 103020 |  | Pacific Blue | BioLegend | IM7 |
| 553133 |  | FITC | BD Biosciences | IM7 |
| 564574 | CD45.1 | BUV737 | BD Biosciences | A20 |
| 110716 |  | APC Cy7 | Biolegend | A20 |
| 110728 |  | PerCP Cy5.5 | Biolegend | A20 |
| 560696 | CD45.2 | PE Cy7 | BD Biosciences | 104 |
| 564616 |  | BUV395 | BD Biosciences | 104 |
| 109824 |  | APC Cy7 | BioLegend | 104 |
| 564449 | CD45R/B220 | BUV737 | BD Biosciences | RA3-6B2 |
| 104438 | CD62L | BV605 | BioLegend | MEL-14 |
| 560513 | CD62L | PerCP Cy5.5 | BD Biosciences | MEL-14 |
| 104508 | CD69 | PE | BioLegend | H1.2F3 |
| 557653 | CD95 | PE Cy7 | BD Biosciences | Jo2 |
| 115803 | CD126 (IL-6Ra) | Biotin | BioLegend | D7715A7 |
| 115806 |  | PE | BioLegend | D7715A7 |
| 25-1271-82 | CD127 (IL-7Ra) | PE Cy7 | Thermo Fischer | eBR2a |
| 13-1321-82 | CD132 (Yc) | Biotin | Thermo Fischer | 10H5 |
| 17-1522-80 | CD152 (CTLA-4) | APC | Thermo Fischer | UC10-4B9 |
| 129804 | CD196 (CCR6) | PE | BioLegend | 29-2L17 |
| 129816 |  | PE Cy7 | BioLegend | 29-2L17 |
| 112705 | CD210 (IL-10R) | PE | BioLegend | 1b1.3a |
| 153504 | CD215 | PE | BioLegend | 6B4C88 |
| 153506 |  | APC | BioLegend | 6B4C88 |
| 12-9949-81 | CD278 (ICOS) | PE | Thermo Fischer | C398.4A |
| 313529 |  | APC/Cy7 | BioLegend | C398.4A |
| 313548 |  | BV711 | BioLegend | C398.4A |
| 25-9985-82 | CD279 (PD1) | PE Cy7 | Thermo Fischer | J43 |
| 551892 |  | PE | BD Biosciences | J43 |
| 13-1219-81 | CD360 (IL-21R) | Biotin | Thermo Fischer | eBio4A9 |
| 633511 | Bcl-2 | PE Cy7 | BioLegend | BCL/10C4 |
| 149029 | CX3CR1 | BV785 | BioLegend | SA011F11 |
| 126406 | FoxP3 | AF488 | BioLegend | MF-14 |
| 25-5773-82 |  | PE/Cy7 | Thermo Fischer | FJK-16s |
| 48-5773-80 |  | eF450 | Thermo Fischer | FJK-16s |
| 565477 | KLRG1 | BV786 | BD Biosciences | 2F1 |
| 557359 | Ly-6C | Biotin | BD Biosciences | AL-21 |
| 560601 | Ly-6G | PE Cy7 | BD Biosciences | 1A8 |
| 108738 | NK1.1 | BV510 | BioLegend | PK136 |
| 740276 | NKG2A/C/E | BUV395 | BD Biosciences | 20d5 |
| 130212 | NKG2D | APC | BioLegend | CX5 |
| 130208 | NKG2D | PE | BioLegend | CX5 |
| 405229 | Streptavidin | BV605 | BioLegend | N/A |
| 644814 | T-bet | APC | BioLegend | 4B10 |
| 109220 | TCRb | APC Cy7 | BioLegend | H57-597 |
| 109227 | TCRb | PerCP Cy5.5 | BioLegend | H57-597 |

#### Single-cell RNA sequencing using the 10X platform

Human or mouse T cells were bulk sorted by FACS into Eppendorf tubes containing cold sterile PBS 10% FCS and incubated for 20 min at 4°C with TotalSeq^TM^DNA-barcoded anti-human or anti-mouse ‘Hashing’ antibodies (BioLegend) at a 1/100 final dilution. The TotalSeq-A^TM^antibodies conjugated to DNA oligonucleotides stained all leukocytes in the relevant samples. During the incubation, cells were transferred into a 96-well round bottom plate, on ice. Following incubation, cells were washed three times in cold PBS 2% FCS and the hashed populations pooled into mixtures for single-cell RNA sequencing using the 10X Genomics platform. In the case of human T cells, the hashed mixtures were first spiked with 3T3HEK mouse cells (2%) and incubated with DNA-barcoded antibodies (see Key Resources Table), to allow high-throughput measurements of cell-surface proteins integrated with transcriptome measurements, by cellular indexing of transcriptomes and epitopes by sequencing (CITE-seq) (Stoeckius et al., 2017).

The Garvan-Weizmann Centre for Cellular Genomics (GWCCG) performed the 10X capture, and sequencing of resulting cDNA samples, as an in-house commercial service, using the Chromium Single-Cell v2 3’ Kits (10X Genomics) as per the manufacturer protocol. A total of 5,000 to 20,000 cells were captured per reaction.

RNA libraries were sequenced on an Illumina NovaSeq 6000 (NovaSeq Control Software v 1.6.0 / Real Time Analysis v3.4.4) using a NovaSeq S4 230 cycles kit (Illumina, 20447086) as follows: 28bp (Read 1), 91bp (Read 2) and 8bp (Index). HASHing libraries were sequenced on an Illumina NextSeq 500/550 (NextSeq Control Software v 2.2.0.4 / Real Time Analysis 2.4.11) using a NextSeq 60 cycles kit (Illumina, 20456719) as follows: 28bp (Read 1), 24bp (Read 2) and 8bp (Index). Sequencing generated raw data files in binary base call (BCL) format. These files were demultiplexed and converted to FASTQ using Illumina Conversion Software (bcl2fastq v2.19.0.316). Alignment, filtering, barcode counting and UMI counting were performed using the Cell Ranger Single Cell Software v3.1.0 (10X Genomics). Reads were aligned to the GRCh38 human (release 93) or mm10-3.0.0 (release 84) mouse reference genomes. Raw count matrices were exported and filtered using the EmptyDrops package in R (Lun et al., 2019).

#### DNA-barcoded anti-mouse Hashing antibodies

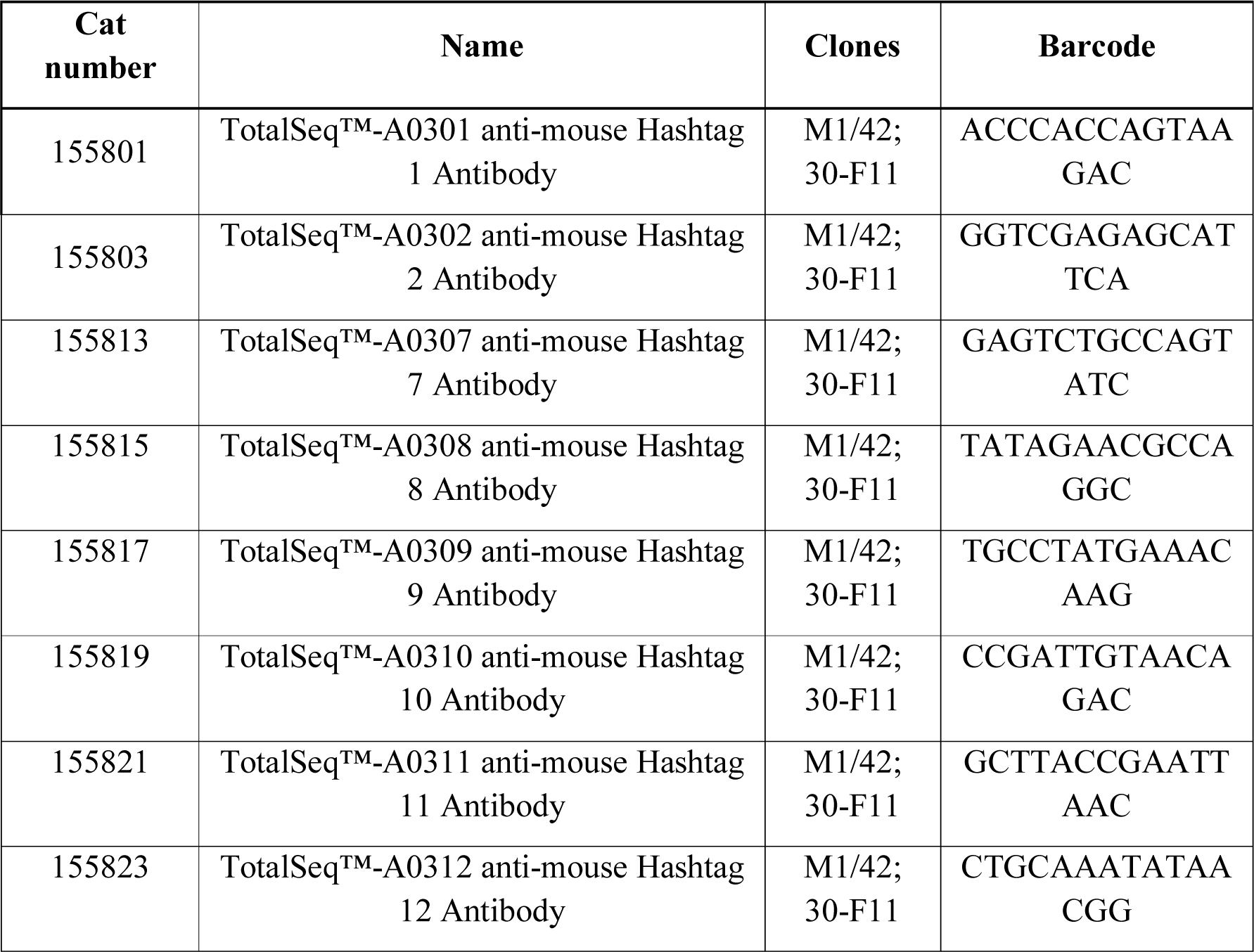

#### DNA-barcoded anti-human Hashing antibodies

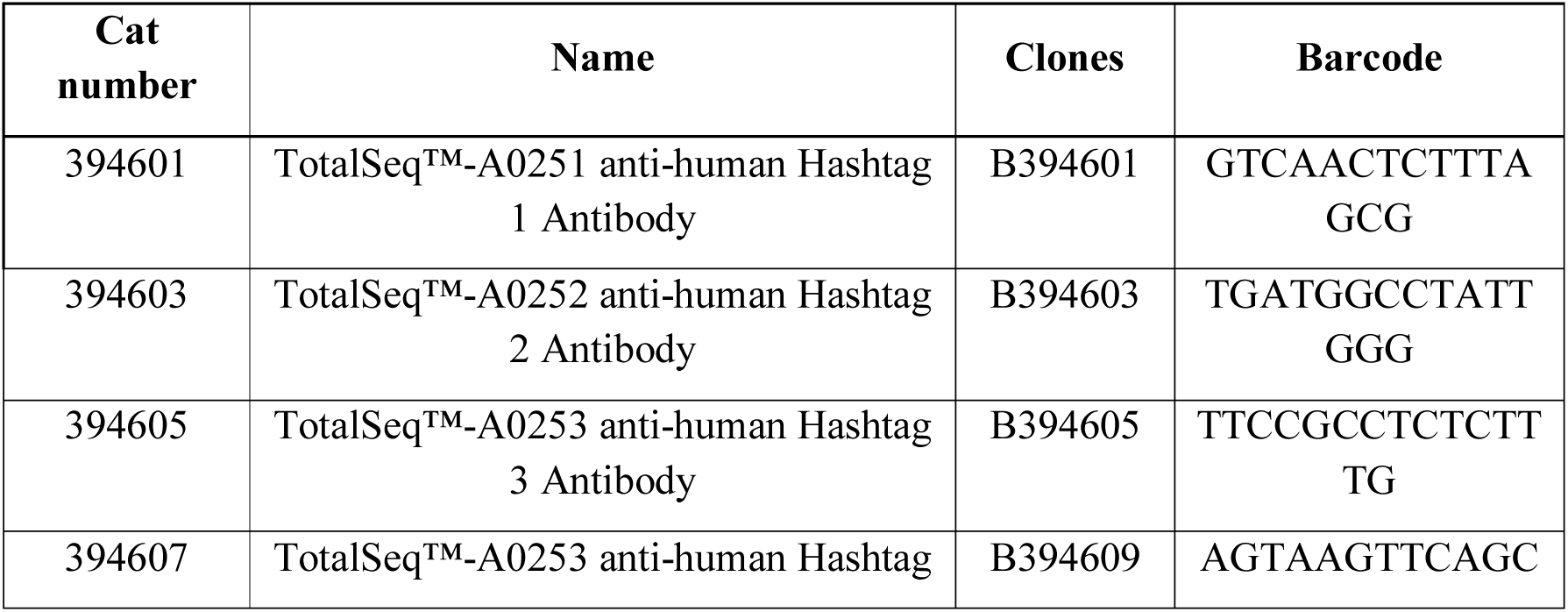
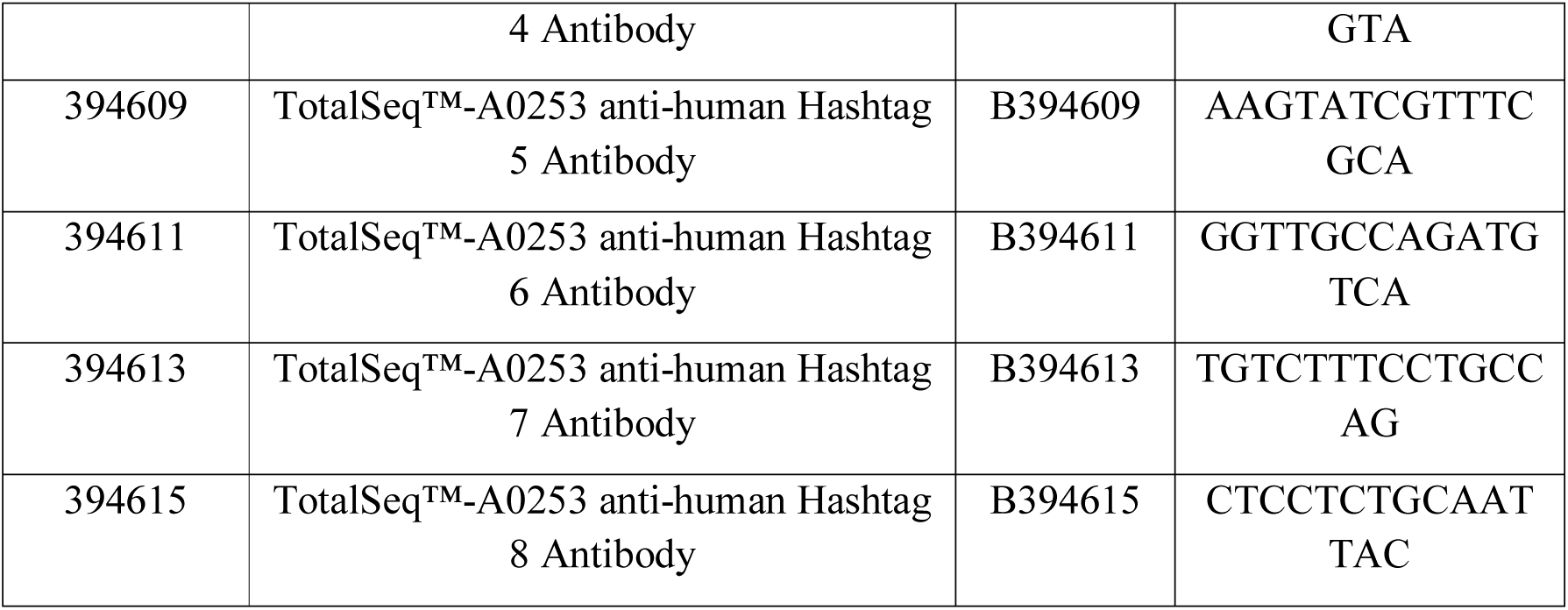

#### Repertoire and Gene Expression by Sequencing

The Repertoire and Gene Expression by Sequencing (RAGE-Seq) method (Singh et al., 2019) was applied in parallel, during 10X sequencing of T cells sorted from healthy controls or individuals with STAT3 GOF syndrome. Briefly, following full-length cDNA 10X capture generation, the cDNA library was split into two prior to fragmentation for short-read sequencing. In parallel to the standard 10X Illumina library preparation and short-read sequencing, full-length sequencing was performed using Oxford Nanopore on selectively-enriched BCR and TCR cDNA transcripts obtained by targeted hybridisation capture, to obtain both the 3’ cell-barcode and the 5’ V(D)J sequence. Cell barcodes were then matched between short-read sequencing and antigen-receptor sequences.

#### TCR deep sequencing by modified 5’RACE

RNA was extracted from T cell populations bulk-sorted by FACS, using the Qiagen AllPrep DNA/RNA Mini Kit (#80204). cDNA synthesis was performed on 2.5 μL of RNA using the cDNA synthesis and library amplification method from the Smart-Seq2 protocol (Picelli et al., 2014). 10 cycles of PCR amplification were performed and the TSO oligo was modified to incorporate a 10 bp unique molecular identified (UMI). Following purification of PCR products with magnetic AMPure XP beads (Agencourt), TCRβ-specific PCR was performed with a primer targeting the constant region of the TCRβ chain and a forward primer (ADP_fwd) targeting the 5’ incorporated TSO oligo.

PCR was performed using the KAPA HiFi HotStart Ready Mix (Kappa Biosystems), under the following conditions: 98 °C 45s; 30 cycles: 98 °C 15s, 60 °C 30s, 72 °C 30s; 72 °C 1min. The PCR products were purified using AMPure beads and complete adaptor sequences and sample barcodes were added using the primers from the Illumina Nextera Index Kit (Illumina). 5 cycles of PCR amplification were performed using the Q5 High-Fidelity DNA Polymerase kit (New England BioLabs): 72 3min; 98 °C 30s; 5 cycles: 98 °C 10s, 63 °C 30s, 70 °C 3min.

Following a second round of magnetic bead purification, the TCR libraries were quantified using the Qubit 4 fluorometer (Invitrogen) and pooled at equal concentration for sequencing on an Illumina MiSeq 250 bp paired-end run.

Sequencing was performed on the Illumina MiSeq platform using the MiSeq Reagent Kit v3 with a read length mode of 2 x 300bp. Libraries were sequenced to ∼1 million reads per sample.

## QUANTIFICATION AND STATISTICAL ANALYSIS

Following flow cytometric experiments, statistical analyses were performed using the GraphPad Prism 6 software (GraphPad, San Diego, USA). A one-tailed unpaired Student’s *t*-test with Welch’s correction was used for comparisons between two normally distributed groups. An unpaired student’s *t*-test, corrected for multiple comparisons using the Holm-Sidak method, was used for comparisons of more than two groups. Differences between paired measurements were analysed by paired *t*-test. In all graphs presented, the error bars represent the mean and standard deviation. * p < 0.05, ** p < 0.01, *** p < 0.001.

Following TCR deep sequencing, sequencing libraries were de-multiplexed by their sample indices using the Illumina FASTQ generation workflow. Paired-end reads were merged using FLASH (Magoc and Salzberg, 2011). Quality filtering, UMI extraction, primer trimming and de-replication were performed using pRESTO (Vander Heiden et al., 2014). The resulting FASTA formatted sequence datasets were aligned against the germline reference directory for the locus and species obtained from IMGT [http://www.imgt.org/] using a local installation of IgBLAST (Ye et al., 2013). IgBLAST alignment determined the V, D and J gene segments that contributed to each rearrangement, any nucleotide insertions or deletions when the segments were joined, the FR and CDRs. TCR repertoires were summarised at the clonotype level, with clonotypes defined as sequences that shared the same V gene, J gene and CDR3 AA sequence.

For nanopore sequencing, RAGE-seq contigs for each cell barcode were generated and TRA and TRB chains for each cell were assigned as described in (Singh et al., 2019). Clonotypes from RAGE-seq were defined and summarised using the same approach as for the repertoire sequencing.

For the 10X analysis of mouse CD8 T cell populations, cells were excluded if the library size or number of expressed genes fell below 2 median absolute deviations, or if mitochondrial reads accounted for more than 20% of total reads. Cell-wise gene expression counts were normalized and recovered using SAVER (Huang et al., 2018) with default values, and differentially expressed genes (DEGs) were identified using limma (Ritchie et al., 2015) on the log-transformed recovered counts. Bonferroni correction was applied to each set of *p*-values. DEGs were defined as having a family-wise error rate (FWER) < 0.05.

For the 10X analysis of human T cell populations, raw BCL files were demultiplexed and aligned to human genome reference GRCh38 and mouse genome reference mm10, using CellRanger v3.0.2 (10X Genomics) to generate UMI count matrices. CellRanger “filtered_barcode” count matrices were processed by removal of any barcodes with less than 100 genes or 1000 UMI’s or with more than 25% mitochondrial content. All cells with between 5-95% mouse UMI’s were classified as having high ambient mRNA (defined as mRNA molecules not present inside a cell before/at time of capture) and therefore removed. R package Seurat (Satija et al., 2015) v3.0.4 was used for normalisation, dimensionality reduction and cluster assignment using default parameters (at resolution 0.8), using all principal components (*n* = 100) found to be significant (*p* ≤ 0.01) as calculated by the Seurat function Jackstraw. Samples were batch-corrected and integrated using R package Harmony v.1.0 using the same statistically significant principal components.

For the CITE-seq analysis of human T cell populations, FASTQ reads were demultiplexed and aligned to the ADT barcode library using package CITE-seq count v1.4.3 ( https://github.com/Hoohm/CITE-seq-Count), using the barcode ‘whitelist’: barcodes meeting quality control requirements following CellRanger v3.0.2 “filter barcode”. UMIs were collapsed using a maximum hamming distance error of 1 for both cell barcode and UMI. The processed ADT UMI count matrices were normalised and scaled using Seurat v3.0.4 default parameters, as per the developer’s protocol for CITE-seq (Stoeckius et al., 2017).

## SUPPLEMENTAL INFORMATION TITLES AND LEGENDS

**Figure S1.**
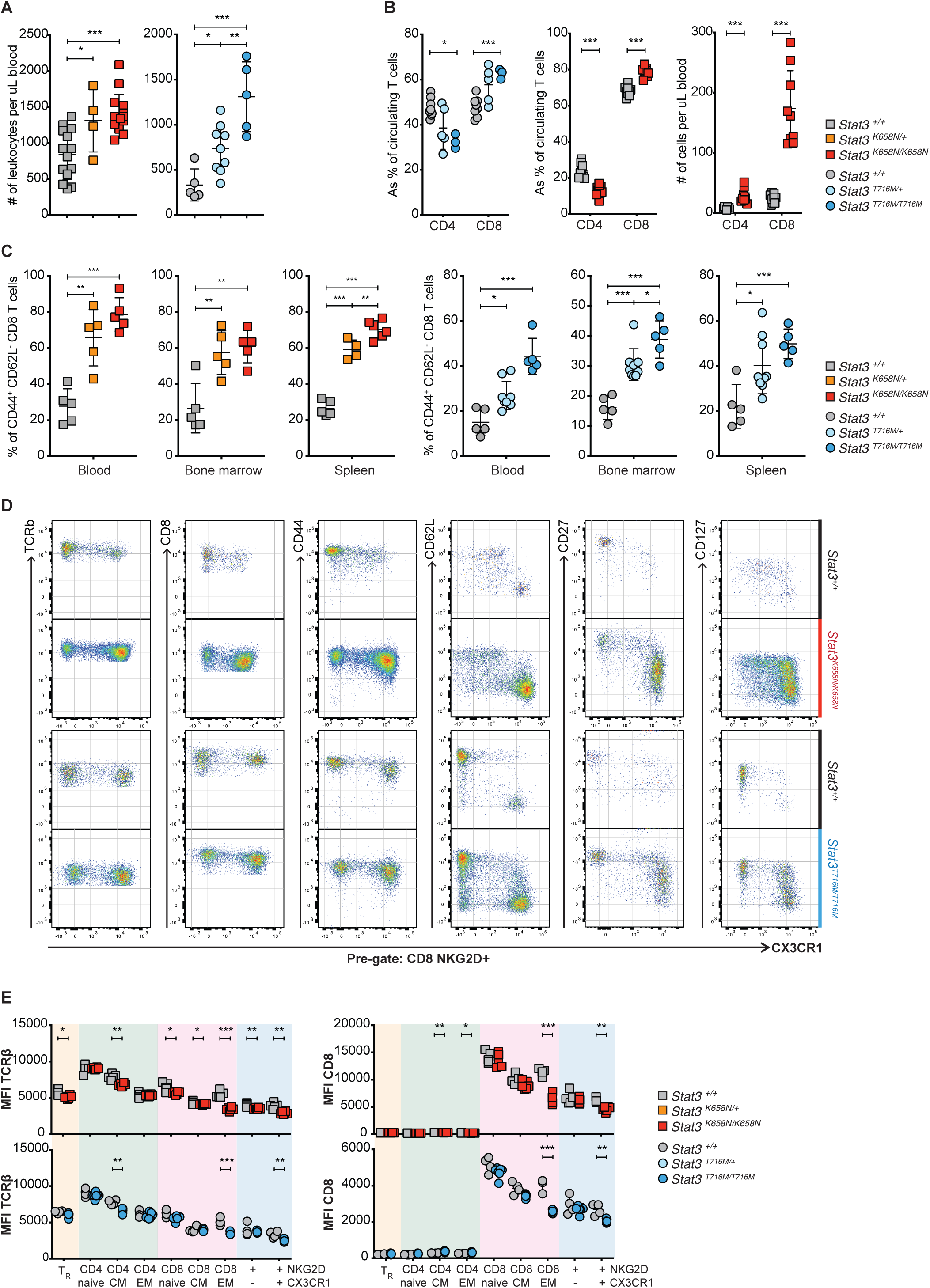
[Germline *STAT3* GOF mutations drive accumulation of NKG2D^+^ CD44^+^ CD62L^-^ CD27^low^CD127^low^ CXCR3^low^CD8 T cells], related to Figure 2. **A.** Symbols show total number of leukocytes per μL blood in individual mice of the genotypes indicated to right of (B). **B.** Percentage of CD4^+^ or CD8^+^ cells among T cells, and number per μL blood in mice of the indicated genotypes. **C.** Percentage of CD44^+^ CD62L^-^ effector memory CD8 T cells that expressed NKG2D in the blood, bone marrow or spleen of individual mice of the indicated genotypes. **D.** Representative flow cytometric analysis of splenic NKG2D^+^ CD8 T cells from mice of the indicated genotypes, showing cell-surface CX3CR1 versus TCRβ, CD8, CD44, CD62L, CD27, CD127. **E.** Mean fluorescence intensity following fluorescent antibody cell-surface staining for TCRβ or CD8, on T regulatory cells (T_R_), naïve, central memory (CM) and effector memory (EM) CD4 and CD8 T cells, NKG2D^+^ CX3CR1^-^ and NKG2D^+^ CX3CR1^+^ CD8 T cells, from *Stat3^K658N^*(top) or *Stat3^T716M^* (bottom) mice of the indicated genotypes. Statistical comparisons made by *t*-test, corrected for multiple comparisons using the Holm-Sidak method. * *p* < 0.05; ** *p* < 0.01; *** *p* < 0.001. For each strain, data are representative of *n* > 3 independent experiments with *n* ≥ 4 mice per group. Data are represented as mean ± SD.

**Figure S2.**
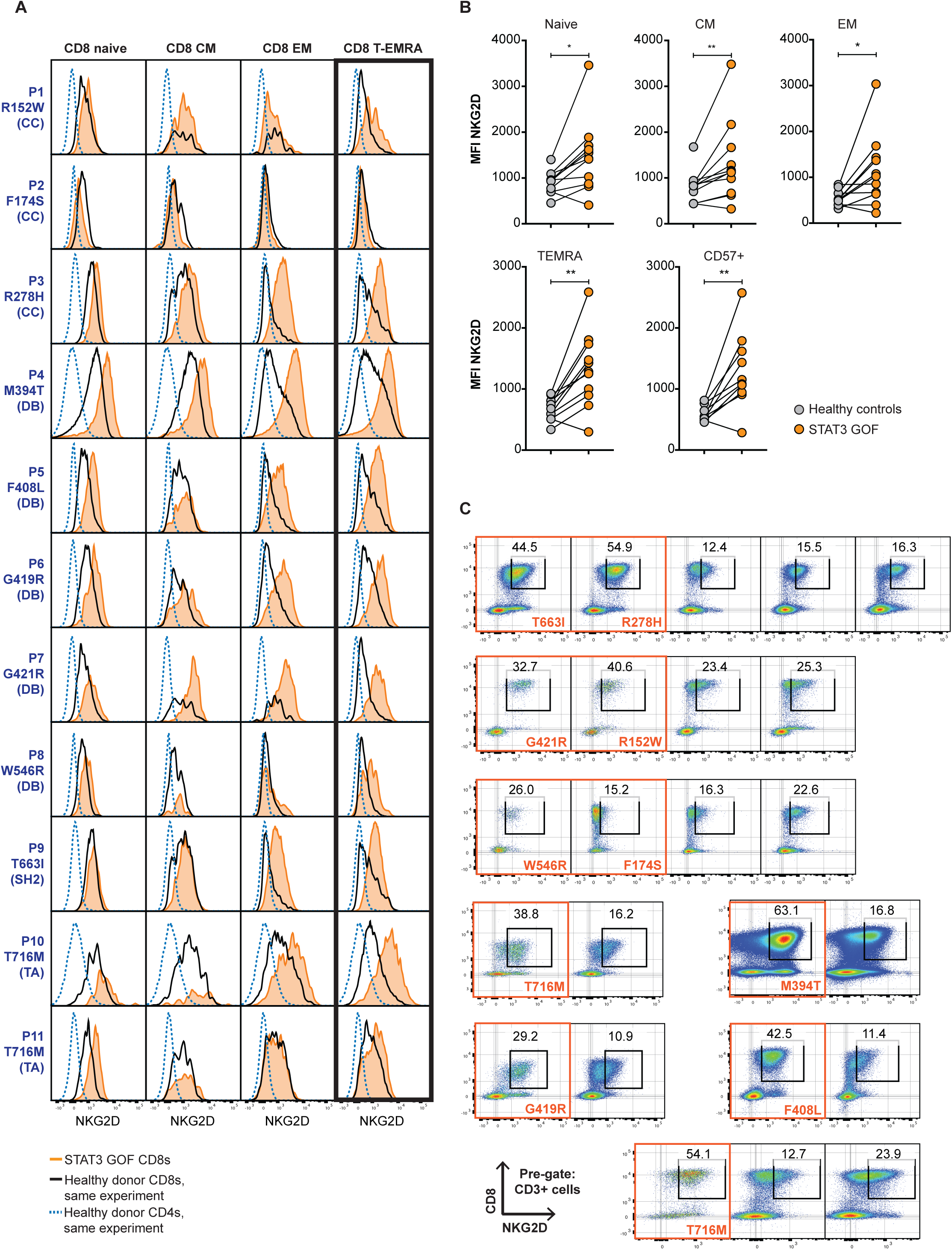
[Increased NKG2D expression and CD8 T cell dysregulation in individuals with STAT3 GOF syndrome caused by mutations modifying different domains of STAT3]**, related to** Figure 3. **A.** Histogram overlays comparing NKG2D cell-surface expression on CD45RA^+^ CCR7^+^ naïve, CD45RA^-^ CCR7^int^central memory (CM), CD45RA^-^ CCR7^-^ effector memory (EM) and CD45RA^+^ CCR7^-^ terminal effector re-expressing CD45RA (T_EMRA_) CD8 T cells from 11 patients with STAT3 GOF syndrome (orange fill), relative to the corresponding CD8 subsets from a healthy donor from the same experiment (black line) and to CD4+ T cells from the same healthy donor (dashed line, serves as negative control as majority of CD4 cells do not express NKG2D). The STAT3 protein domain modified by each patient’s *STAT3* mutation is indicated on the left of the corresponding row. **B.** NKG2D mean fluorescence intensity (MFI) on the indicated CD8 T cell subsets in individual patients and controls. Lines join samples analysed in a same experiment. In experiments that included multiple healthy controls, grey symbols denote the average of the healthy control MFIs. **C.** Plots of NKG2D vs CD8 on CD3+ T cells from each patient and control. Gates show the % NKG2D^hi^CD8+ cells among T cells. Data are combined from *n* = 9 independent experiments each including an equal or greater number of healthy controls to patients. Statistical comparisons made by *t*-test, corrected for multiple comparisons using the Holm-Sidak method. * *p* < 0.05; ** *p* < 0.01; *** *p* < 0.001.

**Figure S3.**
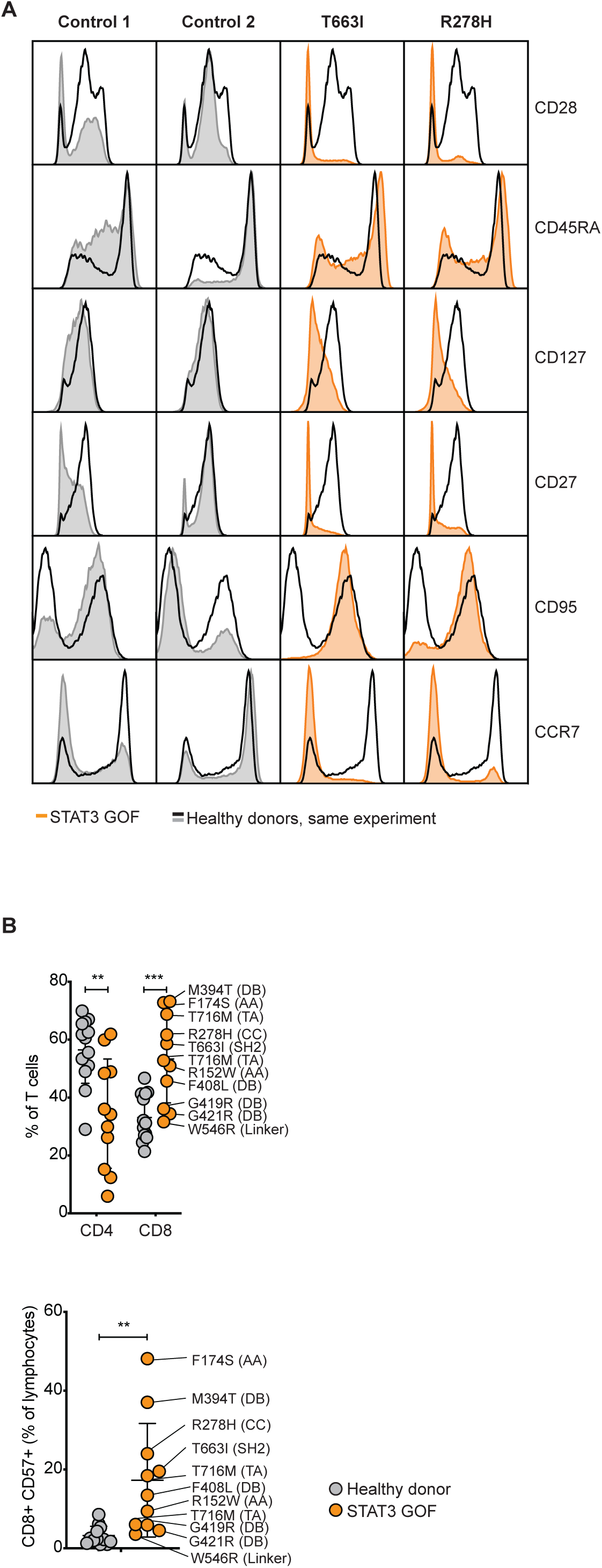
[No discernible correlation between the location of germline mutations in different STAT3 functional domains and the degree of CD57^+^ CD8 T cell dysregulation]**, related to** Figure 3. **A.** Representative histogram overlays comparing CD28, CD45RA, CD127, CD27, CD95, CCR7 cell-surface expression, on CD8 T cells from 2 patients with STAT3 GOF syndrome (orange fill), relative to healthy donors from the same experiment (grey fill, black line). **B.** Percentages of CD4^+^ or CD8^+^ CD3^+^ T cells, and frequencies of NKG2D^hi^or CD57^+^ CD8 T cells as a percentage of total lymphocytes, in the blood of controls (grey) or patients (orange) with corresponding germline STAT3 mutations. Data are represented as mean ± SD. Data are combined from *n* = 9 independent experiments. Statistical comparisons made by *t*-test, corrected for multiple comparisons using the Holm-Sidak method. * *p* < 0.05; ** *p* < 0.01; *** *p* < 0.001.

**Figure S4.**
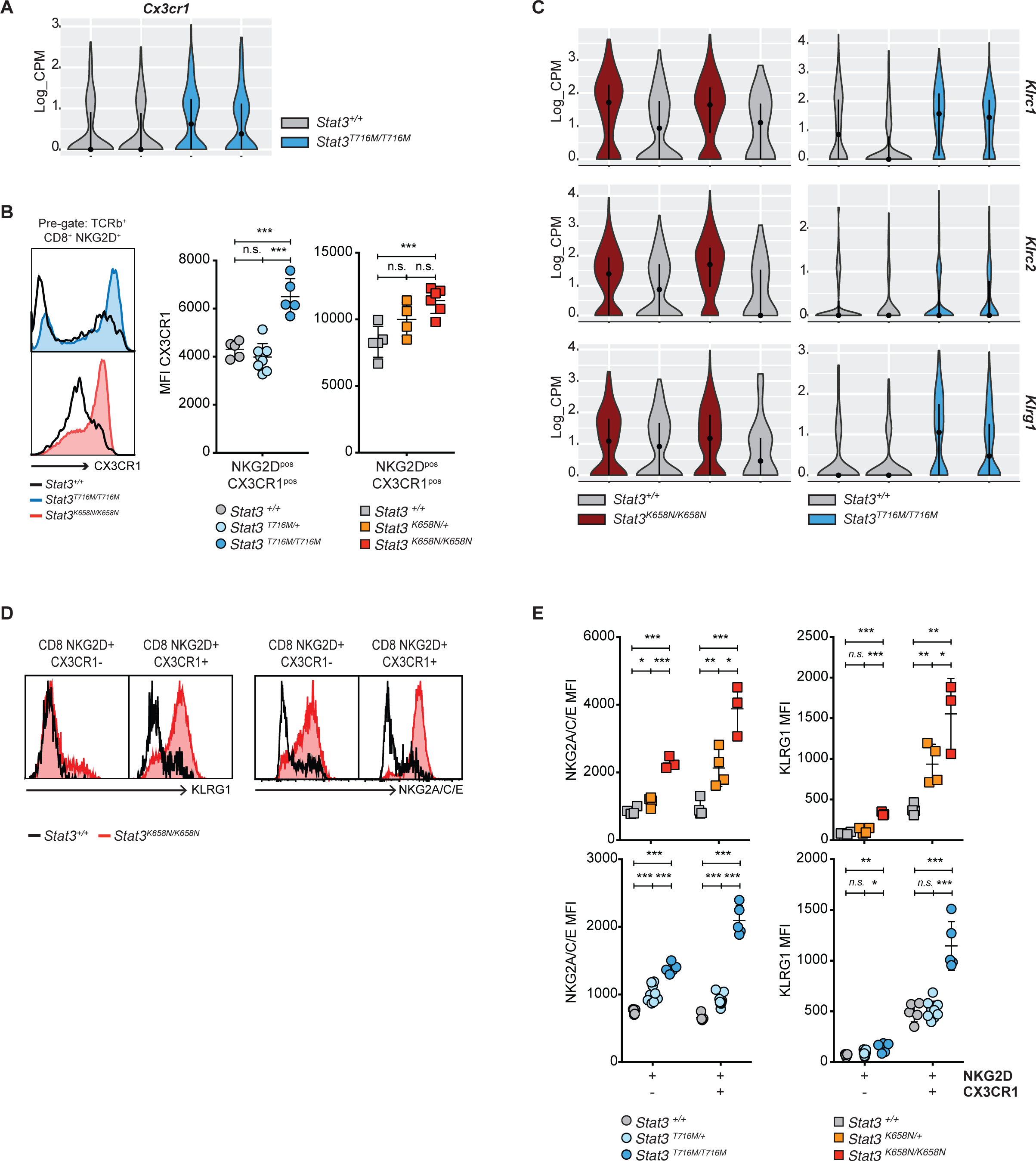
[Validation of increased mRNAs in STAT3 GOF vs wild-type NKG2D^+^ CD8^+^ Cells at the protein level by flow cytometric analysis]**, related to** Figure 5. **A.** Violin plots showing kernel density estimations of *Cx3cr1* gene expression by scRNAseq (log2 normalised counts per million per cell) in single CD8^+^ NKG2D^+^ CX3CR1^+^ T cells from two *Stat3^+/+^*(grey fill) and two *Stat3^T716M/T716M^*mice. Lines denote interquartile range among single cells, and dots geometric mean. **B.** Flow cytometry analysis showing representative histogram overlays of CX3CR1 expression by CD8^+^ NKG2D^+^ T cells in *Stat3^T716M/T716M^*(blue fill; top) or *Stat3^K658N/K658N^*(red fill; bottom) mice relative to *Stat3^+/+^* controls (black line), and mean fluorescence intensity (MFI) for cell-surface CX3CR1 expression by CD8 NKG2D^+^ CX3CR1^+^ T cells from mice of the indicated genotypes. **C.** Violin plots showing kernel density estimations of *Klrc1* (top), *Klrc2* (middle) or *Klrg1* (bottom) gene expression at single-cell resolution, in CD8 NKG2D^+^ CX3CR1^+^ T cells from two *Stat3^+/+^*(grey fill) versus *Stat3^K658N/K658N^*(red fill; left) or two *Stat3^T716M/T716M^*(blue fill; right) mice relative to two littermate *Stat3^+/+^*(grey fill). **D.** Representative histogram overlays of cell-surface KLRG1 (left) or NKG2A/C/E (right) expression by CD8 T cell populations from *Stat3^K658N/K658N^*(red fill) mice relative to *Stat3^+/+^* controls (black line). **E.** Mean fluorescence intensity for cell-surface NKG2A/C/E (left) or KLRG1 (right) expression by CD8 NKG2D^+^ CX3CR1^+/-^ T cells from mice of the indicated genotypes. **B, E.** Statistical comparisons made by *t*-test, corrected for multiple comparisons using the Holm-Sidak method. * *p* < 0.05; ** *p* < 0.01; *** *p* < 0.001. Data are represented as mean ± SD.

**Figure S5.**
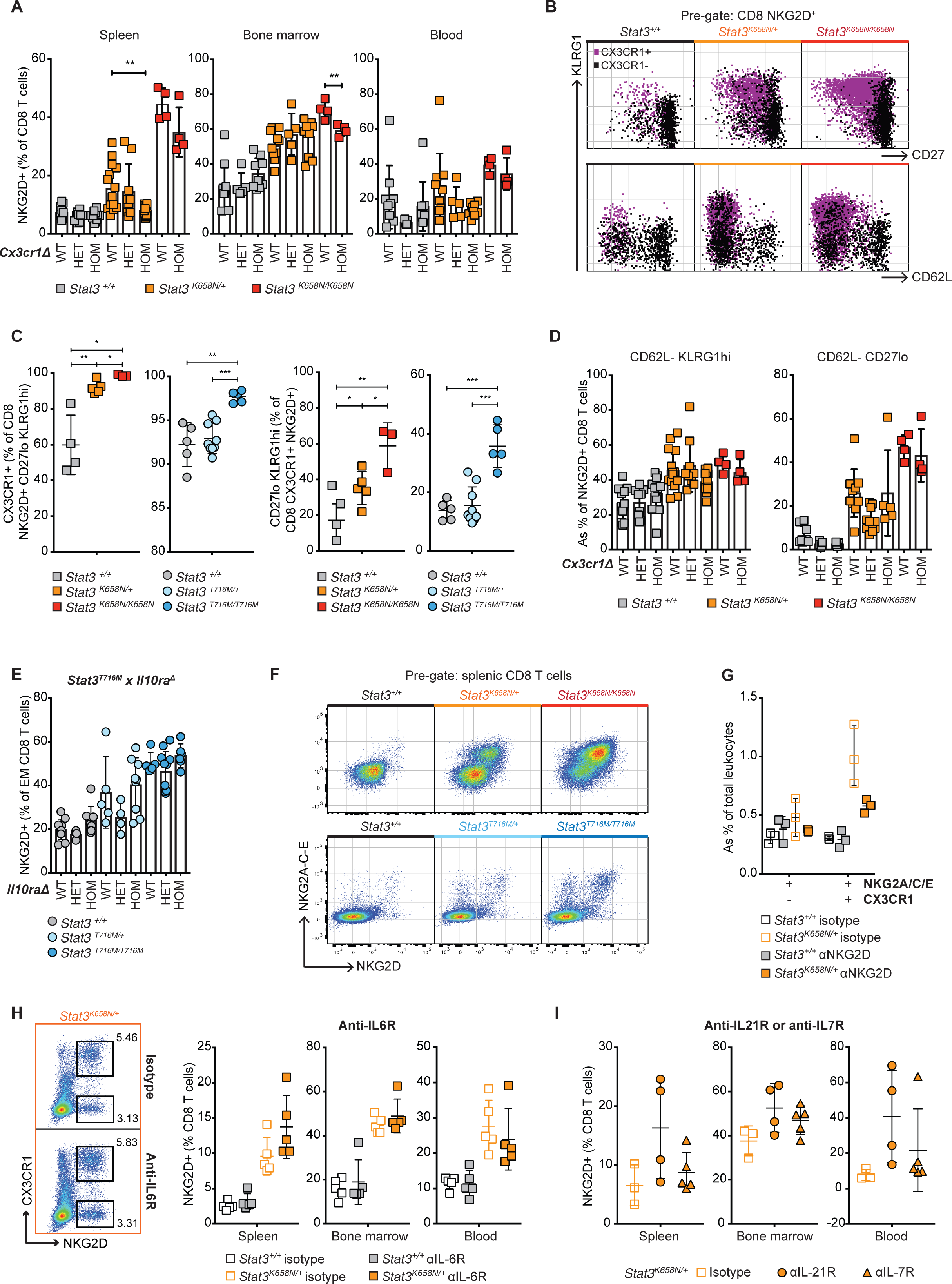
[*Stat3*-mutant CD8 NKG2D^+^ T cells are insensitive to deletion of *Cx3cr1* or *Il10ra*, neutralisation of IL-6R, IL-21R or IL-7R, but sensitive to neutralisation of NKG2D], related to Figure 7. **A.** Percentage of CD8 T cells expressing NKG2D in the spleen (left), bone marrow (middle) or blood (right) of *Stat3^+/+^* (grey), *Stat3^K658N/+^*(orange) or *Stat3^K658N/K658N^*(red) mice that were *Cx3cr1^+/+^*(WT), *Cx3cr1^KO/+^*(HET) or *Cx3cr1^KO/KO^*(HOM) as indicated. **B.** Representative flow cytometric analysis of CD27 or CD62L versus KLRG1 expression, on CD8^+^ NKG2D^+^ T cells from *Stat3^+/+^*(black), *Stat3^K658N/+^*(orange) or *Stat3^K658N/K658N^*(red) mice. **C.** Percent CX3CR1^+^ among CD8 NKG2D^+^ CD27^low^KLRG1^high^T cells (left) or % CD27^low^KLRG1^high^among CD8^+^ NKG2D^+^ CX3CR1^+^ T cells (right), in mice of the indicated genotypes. **D.** Percentage of CD8 NKG2D^+^ T cells that were CD62L^-^ KLRG1^high^(left) or CD62L^-^ CD27^low^(right), in *Stat3^K658N^*x *Cx3cr1^KO^*mice of the indicated genotypes. **E.** Percentage of NKG2D^+^ Cells among splenic CD8^+^ CD62L^-^ CD44^+^ effector memory T cells in *Stat3^+/+^*, *Stat3^T716M/+^*or *Stat3^T716M/T716M^* mice that were *Il10ra^+/+^*, *Il10ra^KO/+^*or *Il10ra^KO/KO^*. **F.** Representative flow cytometric analysis of NKG2D versus NKG2A/C/E expression by CD8 T cells from *Stat3^K658N^*(top) or *Stat3^T716M^*(bottom) wild-type and mutant mice. **G-I.** CD8 T cell subpopulation frequencies in mice that have received 8 bi-weekly injections of neutralising monoclonal antibody (mAb; filled squares/triangles) or isotype controls (empty squares). **G.** Percentage CD8 NKG2A/C/E^+^ CX3CR1^+^ T cells of total splenic leukocytes in *Stat3^+/+^*or *Stat3^K658N/+^*mice treated with anti-NKG2D mAb or control. **H.** Representative flow cytometric plot of NKG2D versus CX3CR1 expression by CD8 T cells and percent NKG2D^+^ Cells (with or without CX3CR1) among CD8 T cells in individual mice treated with anti-IL-6R mAb or isotype control. **I.** Percent NKG2D^+^ Cells among CD8 T cells in individual *Stat3^K658N/+^*mice treated with isotype control (squares), anti-IL-21R (circles) or anti-IL-7R (triangles) mAbs. Statistical comparisons made by *t*-test, corrected for multiple comparisons using the Holm-Sidak method. * *p* < 0.05; ** *p* < 0.01; *** *p* < 0.001. Data are represented as mean ± SD.

**Figure S6.**
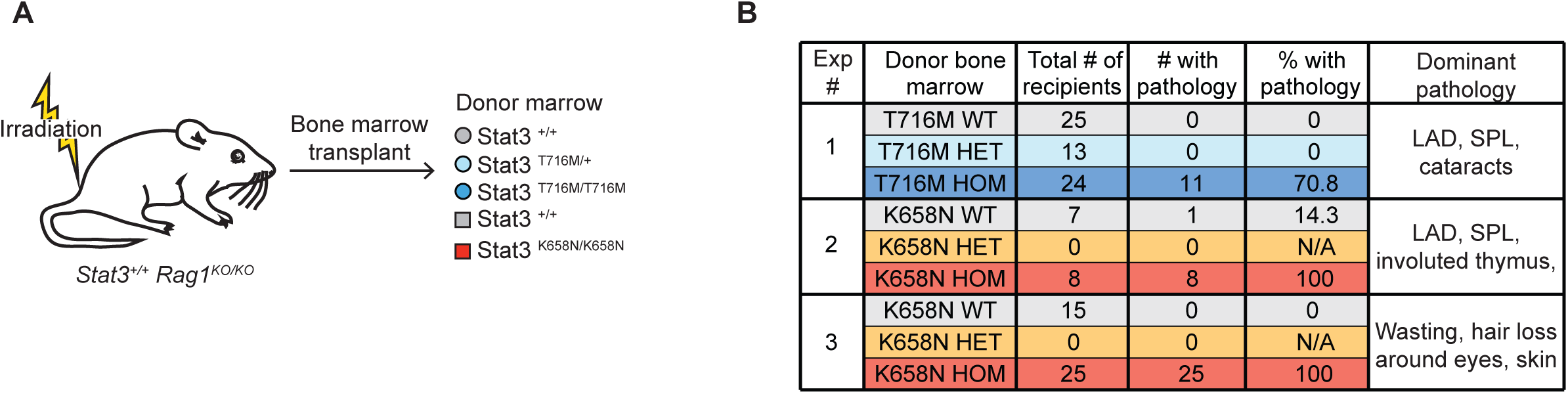
[Transplantation with homozygous GOF *Stat3*-mutant bone marrow causes gross pathology in recipient mice]**, related to** Figure 7. **A.** Experimental design to limit *Stat3* GOF to hematopoietic cells by transplanting mutant bone marrow to *Stat3^+/+^ Rag1^KO/KO^*mice. **B.** Number and % of bone marrow recipient mice developing at least one of the following pathologies: lymphadenopathy (LAD), splenomegaly (SPL), hepatomegaly, cataracts/eye inflammation, skin inflammation on the snout or ears, ringtail, or evidence of gastrointestinal tract inflammation or diarrhea.

**Table S1. [Mutated amino acid residue and STAT3 domain, treatment status, and total white blood cell count from patients with germline or somatic STAT3 mutations], related to** **Figure 3**.

**Table S2. [Differential gene expression in STAT3 GOF relative to healthy donor CD8 T cells]**, related to Figure 4. Column A: gene name. Column B: p value. Column C: log2 fold-change of the average expression for each gene in STAT3 GOF CD8 T cells relative to healthy donor CD8 T cells. Columns D&E: the percentage of STAT3 GOF CD8 T cells (D) or healthy donor CD8 T cells (E) expressing each gene. Column F: adjusted p value.

**Table S3. [Differential gene and protein expression between unique clusters identified by unsupervised dimensionality reduction analysis of non-naïve CD8 T cells in healthy donors or individuals with STAT GOF syndrome], related to** **Figure 4**. Column A: gene name. Column B: *p* value. Column C: log2 fold-change of the average expression for each gene in cells from relevant cluster (indicated in column G) relative to cells of other clusters. Columns D&E: the percentage of cells within (D) or outside (E) the relevant cluster that express each gene. Column F: adjusted *p* value. Column G: unique clusters identified by unsupervised dimensionality reduction analysis.

**Table S4. [Genes differentially expressed in Stat3^T716M/T716M^mice, between NKG2D^+^ CX3CR1^+^ CD8 T cells and CD8 T cells excluding NKG2D^+^ CX3CR1^+^ CD8 T cells], related to Figure 5**.

**Table S5. [Genes differentially expressed in *Stat3^T716M/T716M^*NKG2D^+^ CX3CR1^+^ CD8 T cells relative to *Stat3^+/+^*NKG2D^+^ CX3CR1^+^ CD8 T cells]**, related to Figure 5.

**Table S6. [Genes differentially expressed in Stat3^K658N/K658N^NKG2D^+^ CX3CR1^+^ CD8 T cells relative to Stat3^+/+^ NKG2D^+^ CX3CR1^+^ CD8 T cells], related to Figure 5**.

**Table S7. [Results of gene set enrichment analysis comparing *Stat3^K658N/K658N^*relative to *Stat3^+/+^*NKG2D^+^ CX3CR1^+^ CD8 T cells for sets of genes up-regulated in CD3^+^ CD8^+^ CD44^+^ NKG2D^+^ T cells relative to CD3^+^ CD8^+^ CD44^+^ NKG2D^-^ T cells sorted from lymph nodes of autoimmune alopecic C3H/HeJ mice**], related to Figure 5.

